# Clonal embeddings allow exploratory analysis of lineage-resolved single-cell data

**DOI:** 10.64898/2026.04.30.720820

**Authors:** Sergey Isaev, Alek G Erickson, Igor Adameyko, Peter V Kharchenko

## Abstract

Assays coupling high-throughput lineage tracing with single-cell transcriptomics are transforming studies of development and disease biology, revealing not only major differentiation routes but also continuous fate biases and their putative regulators. Yet, analysis of such data at scale presents challenges due to the sparse nature of clonal data and annotation dependencies. Towards that aim we developed a machine learning approach – clone2vec – which learns informative clone embeddings directly from the cellular expression manifold, bypassing discrete cell-type labels and remaining stable when clones are represented by few cells. This representation summarizes clonal variation as an interpretable geometry that supports exploration, statistics for clone-gene associations, and cross-dataset alignment. In prospective barcoding datasets spanning embryogenesis, tumorigenesis, and hematopoiesis, clone2vec recapitulates established clonal patterns and uncovers new axes of continuous variation that implicate regulatory programs and developmental pathways. In tumor microenvironments profiled with TCR sequencing, clone2vec robustly recovers distinct Treg lineages as well as conserved CD8^+^ T cell sublineages across cancer types, including several bystander-like clonal subsets. Overall, clone2vec provides a robust, general solution for the exploratory analysis of lineage-coupled scRNA-seq data.

---

Arising from a single cell, multicellular organisms develop through clonal expansion and fate specification to create, maintain, and regenerate tissues. The gradual diversification allows initially identical progenitors to explore vast phenotypic spaces. Consequently, the value of exploring clonal variation – whether in morphogenesis, stem cell niche dynamics, or immune responses – lies in its ability to reveal the specific intrinsic and extrinsic factors that shape variability in lineage behavior. Lineage variability is not merely developmental noise, but a functional property of living systems. It reflects how individual progenitors distribute potential across space and time, maintain tissue continuity under local perturbations, and enable organs to be both reproducible and flexible. Clonal diversity is also important for functional and developmental tissue robustness. It is obviously central to the function of the adaptive immune system. In development, clonal heterogeneity can buffer perturbations by allowing some clones to compensate for local losses^1,2^, expand under stress^3,4^, or adjust outputs in response to neighboring deficits^5,6^. Similar clones may diverge in different anatomical or signaling environments, while distinct clones may converge within the same niche^7,8^. Clonal analysis can therefore help to disentangle the interplay between ancestry and position, especially in migratory and multipotent systems shaped by changing local contexts.

Indeed, in developmental biology, lineage tracing can reveal spatial regulatory boundaries invisible to standard anatomical assessment. For example, in the murine hair follicle, tracing has shown that a stem cell’s initial spatial position – in the niche bulge or hair germ – is the primary predictor of its fate^9^, with local niche cues actively reprogramming cells to stem-like or differentiated states. Similarly, in vertebrate limb mesenchyme, genetic labeling uncovered a sharp dorsoventral lineage restriction plane that does not coincide with morphological boundaries^10^, indicating that early positional signals imprint progenitors with regional identities that are maintained in the subsequent clonal expansion. Clonal analysis has also revealed many contexts in which fate choice is strongly influenced by cell-intrinsic priming, often by exploiting the shared ancestry of sister cells. In the hematopoietic system, for example, lineage tracing has demonstrated that stem cells (HSCs) belonging to the same clone exhibit highly stereotypic behaviors, including sustained lineage bias and specific stress responses^11^. This cell-intrinsic programming is sufficiently stable that clonally related cells maintain their distinctive biases even when separated and transplanted into independent recipients, with their clonal biases overriding the influence of local microenvironments^11,12^. As well, lineage tracing also allows researchers to identify cases when this intrinsic similarity is overridden by extrinsic signals. In the CD8^+^ T cell compartment, sisters derived from a single naive progenitor can diverge dramatically, generating both short-lived effectors and long-lived memory cells, driven by asymmetric exposure to antigen or polarity cues during the initial division^13–16^. Finally, the similarity of clonally related sister cells has also been used to overcome the fundamental limitation of the destructive genomic assays – their inability to measure the same cell at multiple points in time. By using clonally-traced sister cells as proxies for earlier molecular states, researchers have retrospectively approximated cellular trajectories during complex transitions such as *in vitro* reprogramming and drug resistance evolution^17–20^.

To capture clonal dynamics, lineage tracing techniques have evolved significantly from early dyes and single-gene reporters. The introduction of multicolor recombinase systems, such as Brainbow, expanded the resolution of in vivo tracing^21,22^, though spectral overlap limited the number of distinct clones that could be reliably identified. This limitation was overcome by high-throughput genetic barcoding schemes using site-specific recombination or viral libraries, which allow for the tracking of thousands of clones to estimate size distributions and lineage biases^23,24^. More recently, CRISPR “scarring” techniques have enabled the reconstruction of complex lineage trees by accumulating dynamic edits over time, providing an organism-scale view of clonal heterogeneity^8,25^. Crucially, the combination of these technologies with single-cell transcriptome sequencing now permits the simultaneous profiling of lineage history and cell state (e.g. scRNA-seq), enabling the identification of gene-regulatory programs underlying clonal effects.

A number of computational tools has been developed to interrogate lineage-coupled scRNA-seq data in different ways. These include, for example, methods for tree reconstruction^26,27^, inference of early progenitor biasing^28,29^, or lineage-informed cell clustering^30^. A common initial step in analysis of such data, however, is to characterize and navigate the overall diversity of the observed clonal behaviors. This crucial step presents its own computational challenges. Most clones are represented by very few captured cells due to technical undersampling and biological stochasticity, resulting in data sparsity and overdispersion that make the interpretation of individual clonal events unreliable. Heavy-tailed clone-size distributions confound distance measures of clonal similarity. Additional complications arise from convergent edits or homoplasy and barcode collisions in CRISPR-based systems. As a result, clone-level summaries based on cell-type proportions or hierarchical clustering can be unstable and sensitive to the exact cell annotation. Recent methods now yield thousands of clones, making it possible to capture rich clonal variation and probe the underlying developmental logic. However, robust frameworks for large-scale clonal classification and typology – especially in systems where clonal architectures vary dramatically and continually – are still lacking. Exploration and analysis of such data, therefore, necessitates new scalable approaches to quantifying clonal variation and dynamics on the expression manifolds.

Furthermore, a cell type-centric perspective, typically used in exploratory analysis, can fail to capture the full spectrum of clonal variation in the data. First, the resolution and structure of cell type annotations may be insufficient. In the murine in vitro hematopoiesis system^19^, for example, functionally distinct monocyte lineages occupy transcriptionally separate monocyte states that are nevertheless collapsed into a single “monocyte” category when aggregating cells by expression similarity alone (Supplementary Fig. 1a,b). Even at increasing resolution, the annotation boundaries may be ill-defined. In tumor-infiltrating CD8^+^ T cell compartments^31^, for example, phenotypic states such as tissue-resident memory, effector, and transitional populations form continuous gradients rather than discrete clusters (Supplementary Fig. 1c,d). Clustering of such data is likely to produce arbitrary borders, which may distort or obscure clonal variation patterns^32^. A cluster-free approach is therefore needed to quantify similarities between clonal distributions in an unbiased manner.

Here we introduce clone2vec, a robust, annotation-free approach for constructing an effective overview of clonal variation that enables subsequent exploration of recurrent patterns of fate distributions, as well as the analysis of associated molecular features in clonal data. Starting with a continuous representation of expression state, clone2vec embeds clonal data into a metric low dimensional space where clones with similar fate profiles are grouped together. Applied to a variety of clonal tracing datasets, we show that such embeddings provide an informative overview of clonal variation that is robust to sparse sampling. Across embryonic development, human cortical differentiation, and hematopoiesis, clone2vec effectively recovers known clonal patterns while also revealing previously unrecognized substructures. In immuno-oncology datasets spanning diverse tumor types, it resolves a conserved organization of T lymphocyte clones, including multiple bystander populations distinct from tumor-reactive lineages. With its ability to integrate diverse datasets and inferring clone-associated transcriptional and regulatory features, clone2vec offers a practical, general-purpose framework for comparative analysis of clonal dynamics across biological systems.

## Results

### Identification of patterns in sparsely sampled clonal distributions

The analysis of clonal variation generally aims to identify recurrent structured clonal relationships between cells, connecting these patterns to cellular phenotypes and associated molecular features. Assuming that lineage-enabled data generally identifies subsets of cells that originated from the same progenitor cell, we focus on the most basic relationship: similarity of clonal progeny across the cellular expression or state manifold (Fig. 1a and Supplementary Fig. 1f-h). At the limit of dense sampling, progeny can be modeled as distributions and compared using traditional measures such as Wasserstein distance optimal transport (OT) or maximum mean discrepancy (MMD)^33^. However, inferring and comparing such distributions becomes difficult when the sampling is sparse. In most clonal datasets, the distribution of clonal sizes is highly uneven, with most clones being captured by only a handful of cells, while select few are represented by many cells (Supplementary Fig. 1e). This combination of sparsity and imbalance hampers stable estimation of inter-clone similarity, proper identification of which is essential for further data-driven hypothesis generation and testing (Fig. 1b-d).

**Figure 1.**
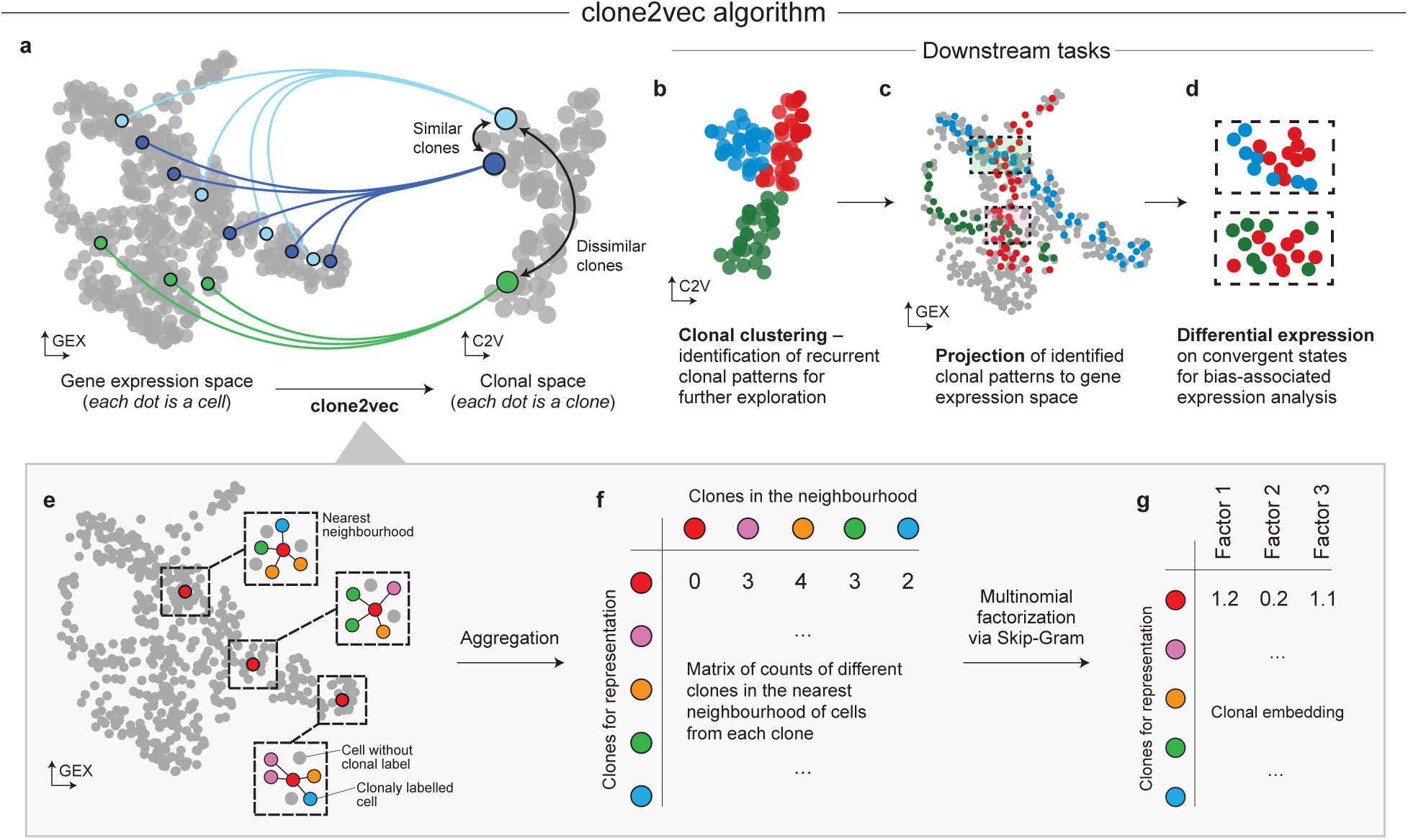
Clonal embeddings with clone2vec. **a**. Schematic of the clonal embedding rationale: clones whose constituent cells occupy similar regions of gene expression space (left) are positioned proximally in the latent clonal space (right), whereas clones with divergent expression distributions are placed distally. Throughout the manuscript, “GEX” denotes embeddings in gene expression space (one point per cell) and “C2V” denotes the clone2vec embedding (one point per clone). **b-d**. Possible downstream analyses enabled by the clonal embedding: (**b**) dimensionality reduction and clustering of the clonal space to identify recurrent clonal patterns; (**c**) projection of clonal cluster assignments back onto the gene expression space to localise convergent transcriptional states; (**d**) differential expression analysis between cells from co-localised but distinct clonal clusters, revealing bias-associated expression programmes. **e-g**. Overview of the clone2vec algorithm. (**e**) For each clonally labelled cell, the k-nearest-neighbour graph in gene expression space is queried (k = 4 shown for illustration). (**f**) Neighbourhood counts are aggregated across all cells of a given clone to yield a clone-by-clone co-occurrence matrix (row for the red clone highlighted). (**g**) The matrix is factorised via a Skip-Gram–based multinomial decomposition to produce a low-dimensional clonal embedding.

To address this problem, we developed clone2vec – a method inspired by the word2vec algorithm for capturing semantic relationships between words^34,35^. Clone2vec first approximates the cellular expression manifold using k-nearest-neighbor graph and uses it to identify which clones co-occur within local neighborhoods (Fig. 1e). It then learns a low-dimensional embedding of clones that best preserves these neighborhood relationships, following a Skip-gram-style objective (Fig. 1f-g). The method can be viewed as a generalized PCA factorization of a neighborhood graph – where exponential family PCA is used to factorize clone-by-clone co-occurrence matrix^36^. Unlike OT or MMD measures that estimate and compare entire empirical distributions, clone2vec aggregates discrete clonal co-occurrence events across expression manifold; we also reasoned that multinomial modelling would yield more robust dimensional reduction under sparse and heavy-tailed clone size distributions^37^.

To evaluate clone2vec’s ability to identify recurrent clonal patterns in sparse sampling regime, we used a synthetic benchmark (Supplementary Fig. 2-4). We simulated distinct patterns (Supplementary Fig. 3a-d) of clonal distributions on a simple manifold, and compared the performance of clone2vec, Skinkhorn divergence, MMD, and cluster-based similarity measures in recovering sets of clones originating from distinct patterns under increasing levels of sparsity (Supplementary Fig. 4a). Indeed, we find that clone2vec shows improved pattern recovery compared to other methods, particularly at high sparsity regimes (Supplementary Fig. 4).

The low-dimensional embedding produced by clone2vec is also analytically convenient, as it supports vector-space algebra. For example, if some clones contribute exclusively to cell type A and others exclusively to B, a clone contributing equally to A and B would be placed at the geometric midpoint between the A- and B-associated clones (Supplementary Fig. 2g-l). Overall, clone2vec embeddings efficiently capture compositional differences between clones in a cluster-free manner, allowing unbiased examination of clonal diversity in the single-cell lineage tracing data.

### Clonal embeddings enable effective exploration of clonal variation patterns

To illustrate how clone2vec summarizes complex clonal variation, we first analyzed central nervous system (CNS) development using data from de Haan and He et al.^38^, where embryos were barcoded by in utero amniotic cavity injection at E7.5 and profiled by scRNA-seq at E9.5 and E10.5. The dataset spans ectoderm-derived neural lineages and includes both progenitor and maturing neuronal populations across the forebrain, midbrain, and hindbrain compartments (Fig. 2a and Supplementary Fig. 5a-c); barcodes enable reconstruction of thousands of multicellular clones captured at these stages. While the main focus of the original work was to describe the relationships between cell types, here we ask a different question: how heterogeneous are the clones and what transcriptional features are linked to this heterogeneity?

**Figure 2.**
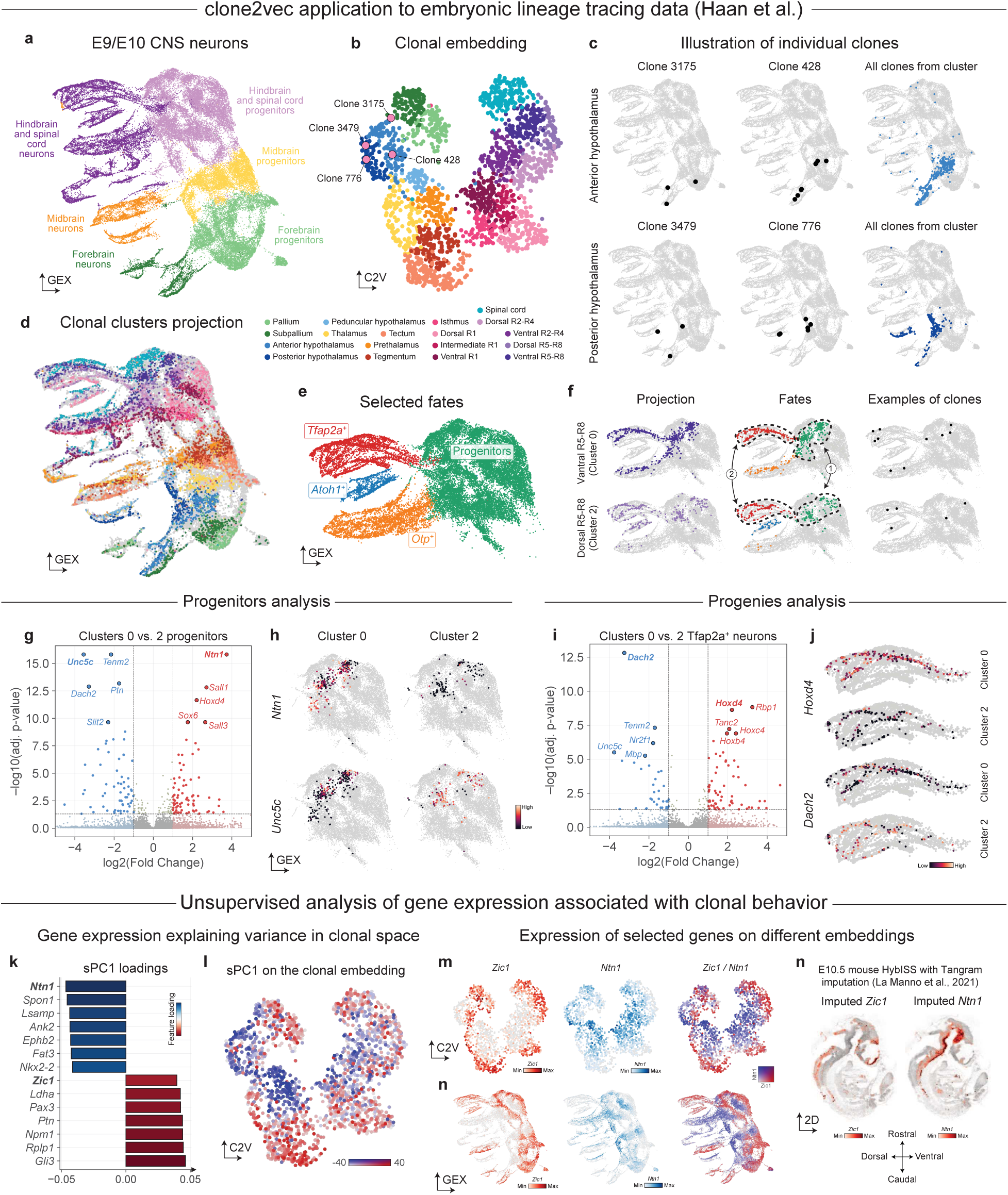
Clonal embeddings reconstruct spatial developmental domains in embryonic data. **a**. Gene expression UMAP of neurons from E9.5/E10.5 murine embryos clonally labeled at E7.5, colored by manually annotated cell types. **b**. UMAP of the clone2vec representation of individual clones, colored by manually annotated clonal clusters. **c**. Gene expression UMAPs showing two example clones (left and center columns) from each of two clonal clusters (rows), and the projection of all cells belonging to clones from these clusters (right column). Hereafter, “projection” refers to all cells from clones of a given clonal cluster shown on the gene expression UMAP. **d**. Gene expression UMAP colored by clonal cluster identity; cells not belonging to clones included in the clonal embedding are shown in grey. **e**. Gene expression UMAP focused on hindbrain and spinal cord neurons and progenitors, colored by manually annotated fine-grained clusters. **f**. Gene expression UMAPs showing the projection (left column) of two clonal clusters (dorsal and ventral R5-R8) onto the focused region, the fine-grained cell-type composition of each projection (center column), and one example clone from each cluster (right column). **g**. Volcano plot of clone-level differential expression in progenitors between ventral (logFC > 0) and dorsal (logFC < 0) populations. P-values: two-sided Welch’s t-test with Benjamini-Hochberg correction, computed on the per-clone average expression across progenitor cells. **h**. Gene expression UMAPs showing expression of *Ntn1* (top row, ventral R5-R8 marker) and *Unc5c* (bottom row, dorsal R5-R8 marker) in cells belonging to ventral (left column) or dorsal (right column) R5-R8 clonal clusters. **i**. Volcano plot of clone-level differential expression in Tfap2a^+^ neurons between ventral (logFC > 0) and dorsal (logFC < 0) populations. Statistics as in (g). **j**. Gene expression UMAPs (single column) showing expression of *Hoxd4* (rows 1-2, ventral R5-R8 marker) and *Dach2* (rows 3-4, dorsal R5-R8 marker) in cells belonging to ventral (rows 1, 3) or dorsal (rows 2, 4) R5-R8 clonal clusters. **k**. Bar plot of loadings of the top 7 genes with positive and negative loadings on the first supervised principal component (sPC1; supervised PCA introduced in Supplementary Fig. 8j-n). **l**. clone2vec UMAP colored by the sPC1 coordinate of each clone. **m**. clone2vec (top row) and gene expression (bottom row) UMAPs showing expression of *Zic1* (left column, high positive sPC1 loading) and *Ntn1* (middle column, high negative sPC1 loading), and their relative expression (right column). **n**. Imputed expression of *Zic1* (left) and *Ntn1* (right) on an E10.5 mouse embryo, obtained from^39^.

Applied to the CNS subset, clone2vec embedding revealed gradual variation in clonal progeny patterns, which we summarized into clonal clusters for interpretation (Fig. 2b and Supplementary Fig. 5d). Visualizing cells belonging to different clonal clusters in cell expression space, we find that individual clonal clusters break up the expression embedding in a manner that would not be apparent from the expression data alone. Most clonal clusters connect specific progenitor subpopulations with one or more differentiating neuronal branches (Fig. 2с and Supplementary Fig. 6). Multiple clonal clusters can contribute to the same neuronal branch (Fig. 2d), and in some regions distinct clonal clusters overlap in expression space indicating minimal expression difference. The top-level geometry of the clonal embedding mirrors the rostral-caudal patterning of the embryonic brain (Supplementary Fig. 5e-h). Interpreting more subtle patterns of clonal variation based on the average expression of cells in each clone, we find that such higher-resolution clonal clusters correspond to well-defined anatomical structures within the developing brain (Fig. 2b,d and Supplementary Fig. 5i). These include telencephalic regions (*Foxg1*^+^), separating pallium with excitatory neurons (*Dmrta2*^+^) and subpallium with inhibitory neurons (*Dlx1*^+^); anterior-posterior hypothalamic territories separated with *Nkx2-1*, *Foxd1* and *Sim1*; or rhombomeric dorsal-ventral sectors (e.g., R1-R4 and R5-R8) resolved with combinations including Hox-genes together with *Il17rd*, *Msx3*, *Nkx6-2*, *Dbx1* and *Lhx1*. Overall, the clonal variation of CNS neuronal progenitors at this developmental stage is consistent with the model in which early clonal diversification is constrained primarily by position within the patterned neural tube.

To illustrate how clone2vec can improve sensitivity of the analysis, we next re-analyzed a recent small-cell lung cancer (SCLC) lineage-tracing study by Ireland et al.^40^ (Supplementary Fig. 7a). The study used lentiviral barcoding of basal-cell-derived organoids and allografts to quantify clonal plasticity in MYC-driven and ASCL1-deficient contexts. The original analysis defined selected clones with >5 cells and used cell-type proportions to define six principal clonal patterns that described clonal plasticity across neuroendocrine, tuft-like, ATOH1, basal and subtype-low states. Applying clone2vec we expanded the set of clones by admitting clones with ≥2 cells, increasing the number of considered clones from 83 to 205, and obtaining robust clone-level embeddings without relying on the cell-type labels (Supplementary Fig. 7b-d). The resulting clone2vec embedding reproduced all of the patterns described by Ireland et al. (Supplementary Fig. 7e), while resolving additional finer distinctions (Supplementary Fig. 7f). Specifically, within the basal-to-ASCL1+ neuroendocrine transition axis (Pattern 1 in Ireland *et al.*), clone2vec resolves three functionally distinct clonal programs (Supplementary Fig. 7g-i). One program is restricted to an epithelial-like SCLC-A state, marked by *Epcam*, *Cd9*, and *Krt8* (cluster 2); a second is restricted to an early neuronal-like SCLC-A state, marked by *Stmn2*, *Dcx*, and *Nrxn3* (cluster 5); and a third spans both programs, consistent with clones that traverse the epithelial-to-neuronal shift (cluster 0). Clone2vec further distinguishes neuronal-like neuroendocrine subtypes: one group displays a *Tshz2* program (cluster 7; corresponding to Pattern 6 in Ireland et al.), whereas the other shows *Zic5* expression, consistent with a neural crest progenitor-like signature (cluster 5). This finer stratification of Pattern 1 suggests additional structure within neuroendocrine plasticity that may be helpful in studies of heterogeneity in drug resistance and state transitions. Overall, clonal embeddings generated by clone2vec provide sensitive representations of clonal variation, enabling further exploration and hypothesis generation on such data.

### Analysis of genes associated with various aspects of clonal variation

A recurring question in the analysis of clonal heterogeneity is how to identify gene expression programs associated with different aspects of clonal behavior. Current analyses have generally focused on targeted tests, typically relating gene expression to the likelihood of a particular cell fate. Such tests can be difficult to interpret when changes involve coordinated shifts among multiple fates. Clonal embeddings provide an explicit representation of this structure and support a more unbiased gene-association analysis.

To illustrate gene expression association testing we first return to the murine CNS development dataset from Haan et al.^38^ As an example, we focus on *Sox1*^+^ neuronal progenitors, which differentiate into three distinct branches: *Tfap2a*^+^, *Atoh1*^+^, and *Otp*^+^ neurons (Fig. 2e and Supplementary Fig. 5j). Clones in cluster 0 span progenitors together with the *Tfap2a*^+^ and *Otp*^+^ branches, whereas clones in cluster 2 span progenitors together with the *Tfap2a*^+^ and *Atoh1*^+^ branches (Fig. 2f). As a first question, we seek to identify gene expression programs in progenitor cells which are associated with these clonal behaviors. To do so, we performed clone-level differential expression between progenitors in clusters 0 and 2 (Fig. 2g-h). Cluster 0 progenitors upregulated *Ntn1*, *Sall1/Sall3*, *Hoxd4*, and *Sox6*, while cluster 2 progenitors upregulated *Unc5c*, *Tenm2*, *Dach2*, *Ptn*, and *Slit2*, consistent with the ventral and dorsal R5-R8 identities of these populations. A complementary question is whether cells of the same terminal fate inherit distinct signatures depending on clonal origin. Analogous differential expression between *Tfap2a*^+^ neurons in clusters 0 and 2 (Fig. 2i-j) recovered posterior Hox genes (*Hoxd4*, *Hoxc4*, *Hoxb4*), *Rbp1*, and *Tanc2* in cluster 0, and *Dach2*, *Tenm2*, *Unc5c*, *Nr2f1*, and *Mbp* in cluster 2. Comparing DE scores across the two tests (Fig. S4k-l) showed that most bias-associated genes are shared between progenitors and their *Tfap2a*^+^ progeny (e.g., *Hoxd4*, *Dach2*), while *Ntn1* and *Sall3* act as progenitor-restricted and *Plxna4* as neuron-restricted markers. As another example of targeted testing we have also re-analyzed traced data on *in vitro* hematopoiesis from Weinreb et al.^19^ (Supplementary Fig. 8). As in the original paper, the analysis focused on comparison of two monocyte-contributing lineages, only one of which contributed to neutrophils. Comparing the two clonal clusters which captured this phenotypic variation, clone2vec re-analysis recapitulated associated expression differences in both progenitor (*Mpo*, *Elane*, *Ctsg*) and differentiated monocyte populations (*Lrg1*, *S100a8*) (Supplementary Fig. 8h,i). In this relatively simple case, the distinction between these two clonal behaviors could be obtained via simple gating on the presence or absence of the neutrophil fate, as was done in the original publication. The advantage of clone2vec embeddings and clonal clusters, however, is that they can capture more complex scenarios, such as those involving changes in multiple cell fates, or situations where cell annotation is not obvious or not sufficiently informative about differences in clonal behaviors.

To extend the expression associations analysis beyond the cluster-level comparisons, we implemented gene-association testing directly on clonal embeddings, using supervised PCA approach (sPCA) in which clone-clone similarities from the embedding define the target kernel, and gene-level association is assessed via contributions to the resulting supervised components^41,42^ (Supplementary Fig. 8j-k). Application of this strategy to the Haan et al. dataset showed the first supervised component associated with clonal embedding to be driven by genes known to be involved in dorso-ventral patterning (*Ntn1*, *Spon1*, *Ank2 vs. Gli3*, *Ptn*, *Pax3*, *Zic1*) (Fig. 2k-l). This was consistent with the spatial transcriptomics data (Fig. 2m-o), demonstrating that the strategy can identify gene programs linked to broad clonal phenotypic structure.

Analogous unsupervised strategy can also be applied to test for the association of clonal phenotypes with gene expression in a particular cell type. Returning to the *in vitro* hematopoeisis example from Weinreb et al., we applied this strategy to examine clonal patterns involving monocytes. We first constructed a new clonal embedding that masked the heterogeneity within the monocyte population itself, by collapsing all monocytes into a single state (Supplementary Fig. 8l). Such clonal embedding is sensitive to overall proportion of monocytes, but not sensitive to their finer state features. We then applied sPCA to identify major expression within monocytes that are associated with the captured clonal variation. The expected neutrophil axis described earlier was captured by the second component, with the top genes matching neutrophil-biasing genes described by Weinreb *et al.* (Supplementary Fig. 8n-p). Surprisingly, the first component was linked to the proportion of undifferentiated cells (Supplementary Fig. 8m) – the axis of variation that could be overseen in direct hypothesis-driven testing.

These analyses show that clonal embeddings can provide a quantitative framework for gene-association testing. This makes it possible to identify gene-expression programs associated with clonal behavior at varying levels of resolution, including multivariate patterns of fate variation that are not readily accessible without a robust numerical representation of fate composition.

### Analysis of clonal variation in large-scale immune datasets

In lymphocyte populations, receptor sequences can serve as endogenous clonal barcodes. The genes encoding both T and B cell receptors (TCRs and BCRs) undergo V(D)J recombination during lymphocyte development generates highly diverse TCR and BCR junctions. Single-cell assays have been optimized to capture the most variable portion of the receptor, typically summarized by the complementarity-determining region 3 (CDR3) sequence^43^. For T cells, this rearranged sequence is generally stable and directly defines a clonal population, while in B cells antigen-driven somatic hypermutation produces a set of related sequences that together constitute a clonal subpopulation.

To illustrate clone2vec for immune-receptor analysis, we first examined clonal structure in peripheral blood mononuclear cells (PBMCs) from healthy adults. Given recent technological advances and the relative ease of sampling, PBMC datasets can now reach millions of cells^44,45^. To support analysis at this scale, we draw on the recent theoretical results showing that the Skip-Gram objective utilized by clone2vec can be effectively approximated by applying Poisson GLM-PCA to the matrix of clone-neighbor co-occurrences^36,37,46^ (see Methods). This formulation relies on the assumption that neighbor counts are generated via a log-linear interaction of latent embeddings, allowing us to recover comparable clonal embeddings with significantly improved computational scalability (Supplementary Fig. 2-4).

Applying clone2vec with Poisson loss to a PBMCs dataset from Sureshchandra et al.^45^ covering almost 2.3 million cells from 10 healthy donors (Supplementary Fig. 9a), of which 208,683 cells are part of 23,362 clones of size three and more (Supplementary Fig. 9b), we found four main clonal clusters corresponding to two general lineages of CD8^+^ T cells (MAIT and memory) as well as two main lineages of CD4^+^ T cells (memory and regulatory) (Fig. 3a-b and Supplementary Fig. 9c-d). Unlike developmental or differentiation settings, where clones often span multiple cell types, from progenitor to effector types, PBMC clonal clusters are largely restricted to individual cell types. This is expected because the receptor rearrangement that “barcodes” the cell typically occurs alongside lineage commitment, and subsequent inter-lineage transitions are rare^47^. Nonetheless, as we show in the next two sections, receptor-defined clonal structure can uncover functionally critical variations within T lymphocytes, distinguishing the developmental origins of T cell populations, or linking transcriptional states to their history of antigen-driven activation and selection.

**Figure 3.**
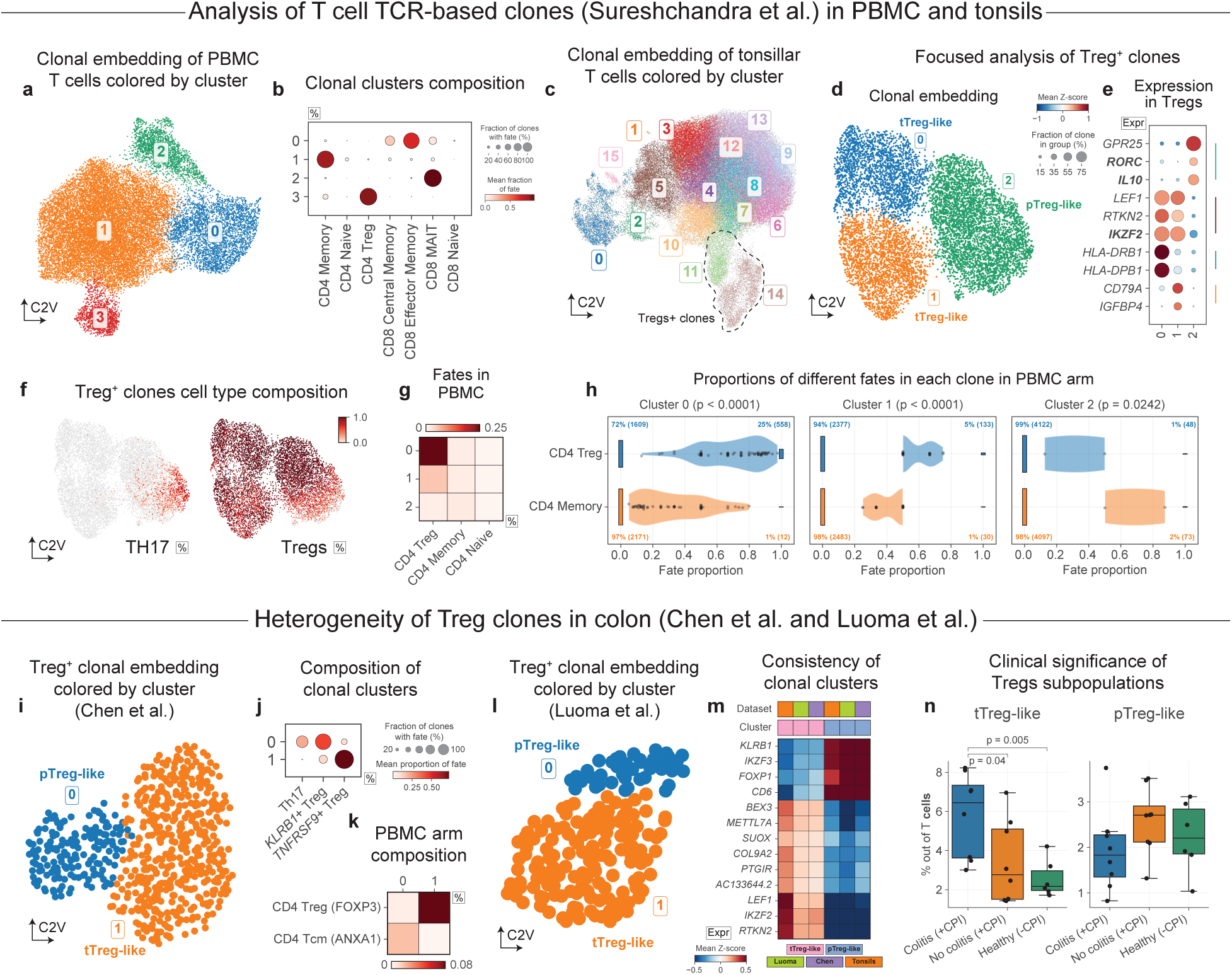
Clonal embeddings capture tTreg- and pTreg-like lineages in CD4 T cells. **a**. clone2vec UMAP of TCR-defined T cell clones from PBMCs of 10 donors (Sureshchandra et al.), colored by Leiden clusters. **b**. Dot plot of cell-type composition for each clonal cluster. Hereafter, in composition dot plots, dot size represents the number of clones containing that fate, and color represents the average proportion of that fate within those clones. **c**. clone2vec UMAP of TCR-defined T cell clones from tonsils of the same 10 donors, colored by Leiden clusters; Treg-containing clusters are outlined with dotted lines. **d**. clone2vec UMAP of Treg-containing clusters from (c), re-clustered, colored by Leiden clusters. **e**. Dot plot of mean marker expression in Tregs from each clonal cluster. Hereafter, in clonal expression dot plots, dot size represents the number of clones with at least one cell expressing the gene, and color represents the average of the per-clone mean expression values. **f**. clone2vec UMAP of Treg-containing clusters, colored by the per-clone proportion of Th17 cells (left) and Tregs (right). **g**. Heatmap of the average fate composition in PBMCs among clones shared with tonsil clones from each Treg-containing cluster. **h**. Modified violin plots (with bars at 0 and 1 sized according to the fraction of values equal to those numbers) of fate distributions in PBMCs for clones shared with each Treg-containing tonsil cluster. P-values: two-sided Mann-Whitney U-test, without multiple testing correction. **i**. clone2vec UMAP of Treg-containing clusters from the colorectal cancer (CRC) tumor microenvironment (TME; Chen et al.), colored by Leiden clusters. **j**. Composition dot plot of the Treg-containing clusters from (i). **k**. Heatmap of the composition of the matched PBMC part for each clonal cluster from the TME. **l**. clone2vec UMAP of Treg-containing clusters from colon (Luoma et al.), colored by Leiden clusters. **m**. Heatmap of mean expression in Tregs of conserved pTreg and tTreg signatures derived in this study by differential expression analysis (see Methods). **n**. Box plot of the proportion of pTreg- or tTreg-like cells relative to all T cells per sample. Center lines, medians; box limits, 25th-75th percentile (IQR); whiskers, 1.5 × IQR. P-values: two-sided Mann-Whitney U-test, without multiple testing correction. Hereafter, box plots follow the same convention.

### Clonal embeddings help to separate thymic and peripherally induced Tregs

Regulatory T cells (Tregs), essential for providing immunotolerance to self, are increasingly understood to arise through distinct developmental routes: thymus-derived Tregs (tTregs), and peripherally induced Tregs (pTregs) generated from conventional CD4 T cells under tolerogenic conditions^48–50^. Furthermore, the pTreg compartment is intimately linked to the plasticity of the Th17 lineage; during the resolution of inflammation, Th17 effectors can undergo context-dependent reprogramming and acquire a regulatory transcriptional program and suppressive function, including *FOXP3*-associated states^51–53^. Distinguishing thymic versus peripheral origin can be challenging, particularly in human studies, as commonly used markers can reflect activation or tissue context rather than lineage history, complicating efforts to map “origin” onto functional state and clonal architecture^54,55^.

As Tregs-containing clones from PBMC appeared to be quite homogeneous, we decided to examine clonal heterogeneity of Treg-associated lineages within the tonsillar part of the same dataset (Supplementary Fig. 9e), consisting of 105,978 clones of size three and more (covering 702,839 cells). Indeed, tonsillar T cell clones showed much higher level of clonal heterogeneity, mostly related to the CD4 T cells lineages (Fig. 3c and Supplementary Fig. 9f-i). Focused analysis of Treg-containing clones revealed three well-separated clonal subtypes with distinct transcriptional signatures (Fig. 3d-e). Treg clonal cluster 2 was notable for containing Th17 cells alongside a distinct Treg state with high *RORC* and low *IKZF2*, which is consistent with pTreg markers established in murine studies^54,56–59^ (Fig. 3f). Additional evidence supporting distinct origin of the putative pTreg cluster comes from the fate distribution of these clones in peripheral blood. The experimental design of the original study included matching PBMC and tonsil samples from the same donors. In blood, the sister cells of clones from the putative tonsillar pTreg cluster 2, were significantly more likely to be found among memory T cells, rather than Ttregs (Fig. 3g,h). In contrast, the sister cells of the other two tonsillar clonal clusters were mostly found among PBMC Tregs.

As Tregs are abundant in colonic mucosa, we next examined single-cell data on colorectal cancer^60^. Clone2vec embeddings largely consist of clusters with the major annotated T cell types (Supplementary Fig. 10a,b), but within the Treg compartment we consistently observed two well-separated clonal clusters that remained stable upon focused re-embedding (Fig. 3i and Supplementary Fig. 10c-f). Treg clonal cluster 0 was notable for containing Th17 cells (Fig. 3j) alongside a distinct Treg state with high *RORC* and low *IKZF2* (Supplementary Fig. 10g), consistent with our previous observations. Comparison of clonotype sharing with matched PBMCs from the same patients suggested divergent lineage connectivity: while *IKZF2*^+^ tTreg-like cells displayed highly Treg-restricted clone sharing, *RORC*^+^ pTreg-like cells shared clonotypes more broadly with circulating CD4 T cell states, including *ANXA1*^+^ Tcm-like cells (Fig. 3k).

We next asked whether this clonal partitioning of Tregs is conserved in other inflammatory contexts by analyzing an independent colon dataset from checkpoint inhibitor-associated colitis by Luoma et al^61^. Here, clone2vec analysis again separated Tregs into two clonal clusters with a concordant marker program (Fig. 3l-n and Supplementary Fig. 10m-n): an *IKZF3*^+^ pTreg-like group and an *IKZF2*^+^ tTreg-like group. Re-analysis of Treg-only gene expression embedding allowed to assign observed Tregs clonal behaviors to Treg cells clusters in gene expression space (Supplementary Fig. 10o-r), revealing additional markers (*IL10* and *RORC* for pTregs) consistent with previous observations. We did not observe Th17 progeny within the pTreg-like clonal cluster, however, these samples generally lack Th17 cells. Altogether, the recurrent clonal split across datasets, specific clonal proximity to Th17 cells, argues that the *IL10*^+^ *RORC*^+^ cluster corresponds to a pTreg-like compartment linked to Th17-associated differentiation trajectories.

Overall, this example illustrates the power of clonal analysis in capturing distinct origins and plasticity within very similar cell populations. Such distinctions can aid studies of disease mechanisms. For example, in Luoma et al.^61^ data on checkpoint inhibitor-associated colitis, we find that the tTreg-like (*IKZF2*^+^) cells was significantly more abundant among patients experiencing colitis complications, while the pTreg-like (*IL10*^+^ *RORC*^+^) cells showed no significant enrichment (Fig. 3n).

### Archetypal lineages of CD8^+^ T cells in tumor microenvironment

Next, we aimed to illustrate how clonal analysis of immune receptor sequences can resolve clonal structures within an individual cell type that may reflect prior antigen exposure, expansion history, or microenvironmental cues. This is particularly important for tumor microenvironment CD8^+^ T cells, which can occupy similar transcriptional states despite very different histories: recent trafficking *vs.* long-term residency, bystander infiltration vs. tumor-antigen engagement, or acute *vs.* chronic stimulation^62^. As checkpoint blockade therapies aim to amplify effective anti-tumor CD8^+^ T cell immunity, clarifying which clones are likely tumor-reactive and how they differentiate is of immediate therapeutic relevance^63^. We therefore analyzed data on non-small cell lung cancer (NSCLC) by Caushi et al.^31^, which analyzed matched tumor, adjacent normal and peripheral blood samples with both TCR sequencing and orthogonal antigen-specificity assays for representative clonotypes, providing an opportunity to connect clonal structure to antigen experience (Supplementary Fig. 11a-b).

Restricting to conventional CD8^+^ T cells (excluding innate-like CD8^+^ populations such as MAIT cells), we constructed clone2vec embeddings for TCR-defined clones across the cohort from Caushi et al.^31^ (Supplementary Fig. 11c,d). Rather than forming a small number of sharply separated clone “types,” the clone embedding space showed a graded spectrum of behaviors (Supplementary Fig. 11e), consistent with tumor-infiltrating CD8^+^ T spanning multiple activation, residency, and dysfunction programs despite sharing a common cell identity^31,62^. We therefore summarized this structure using archetypal analysis (Fig. 4a-d), yielding four dominant clone-level programs that we refer to by their most discriminative markers: *CXCL13*^High^, *IL7R*^High^, *GZMK*^High^, and *S1PR5*^High^ (Fig. 4e).

**Figure 4.**
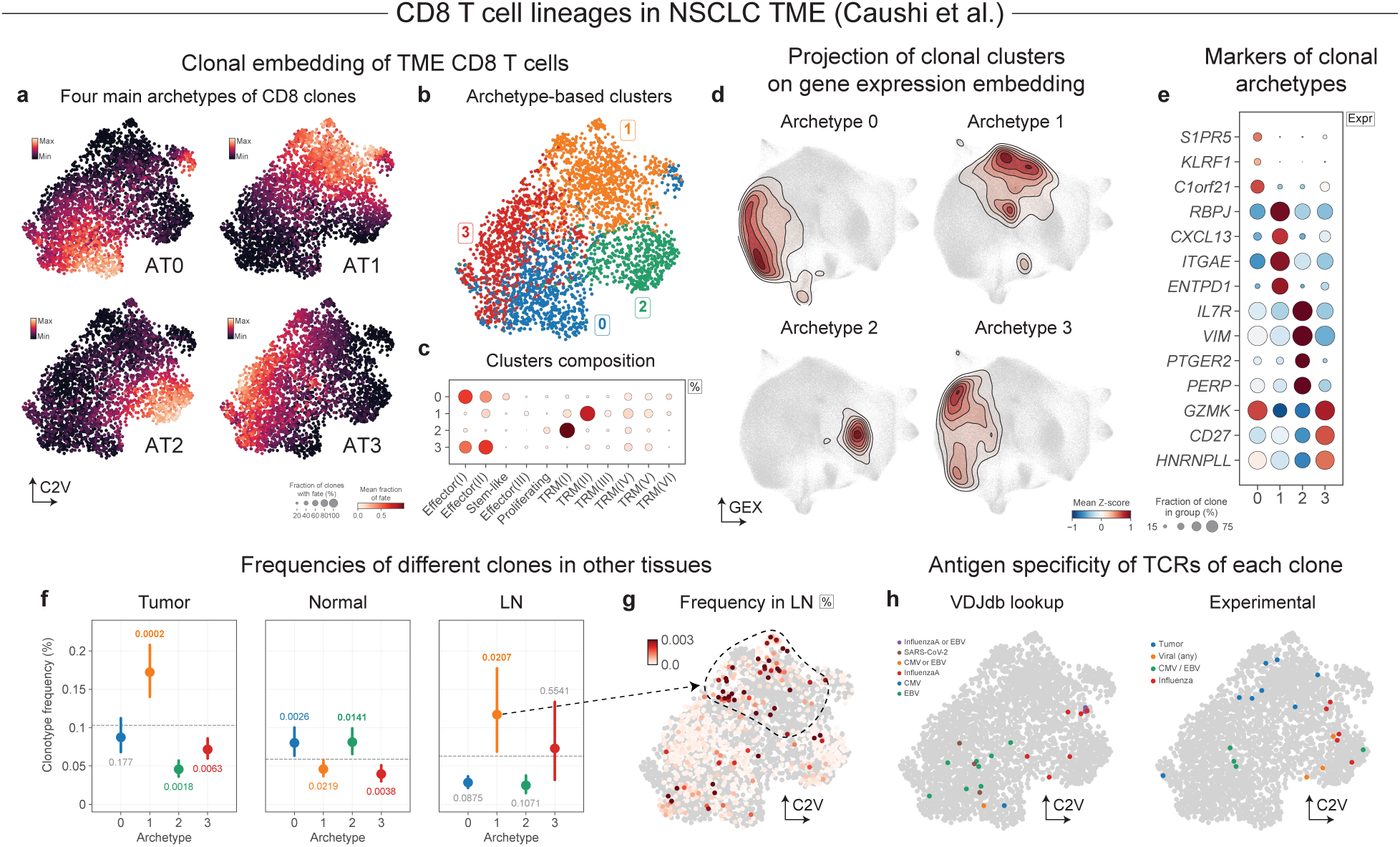
Clonal embeddings reveal four archetypes of CD8 T cell lineages in the lung cancer microenvironment. **a**. clone2vec UMAPs of adaptive-like CD8 T cell clones from the lung cancer microenvironment (Caushi et al.), arranged as a 2×2 grid (top row: archetypes 0, 1; bottom row: archetypes 2, 3), colored by archetype weight. **b**. Same clone2vec UMAP as (a), colored by archetype-based hard clustering, derived as the argmax of archetype weights per clone. **c**. Dot plot of cell-type composition for each archetype-based cluster. **d**. Gene expression UMAPs showing kernel density estimation (KDE) of the projection of each archetype-based cluster. **e**. Dot plot of mean marker expression in CD8 T cells within each archetype-based cluster. **f**. Point plot (mean represented by dots, whiskers showing nonparametric bootstrap-based 95% confidence intervals of the mean with 10,000 bootstrap replicates) of average clonotype frequency for clones from each archetype-based cluster, stratified by tissue. The values above (enriched) or below (depleted) each dot, give p-values for the difference between the cluster’s clonotype frequency and the average across all clusters in that tissue, calculated with a permutation test (100,000 permutations) followed by Benjamini-Hochberg correction. **g**. clone2vec UMAP colored by the average frequency of each clonotype in matched lymph node samples; grey indicates absence of a matched lymph node sample for the donor of the clone. **h**. clone2vec UMAPs colored by VDJdb lookup-based (left) or experimentally identified (right; from the same study, Caushi et al.) antigen affinities.

Interpreting these programs using the study’s orthogonal antigen-specificity measurements and cross-compartment sampling, we found that each archetype likely corresponds to a distinct history of antigen exposure. *CXCL13*^High^ clones expressed a canonical dysfunction signature (including *ENTPD1*/CD39) and accounted for almost all of the TCRs with an experimentally validated affinity to the tumor neoantigen, while lacking other types of validated TCRs. *CXCL13*^High^ clones were also preferentially represented in tumor and draining lymph node regions compared to adjacent normal lung tissue^31^ (Fig. 4f,g and Supplementary Fig. 11f-g). In contrast, *IL7R*^High^ clones were enriched in adjacent normal lung and contained TCRs with affinity to influenza virus (Fig. 4f,h and Supplementary Fig. 11f-g), consistent with resident-memory bystanders in the respiratory tract^31,64^. *GZMK*^High^ clones showed similar distribution across samples, but were enriched for TCRs specific to EBV and CMV (based on both experimental affinity measurements and VDJdb^65^) (Fig. 4f-h), consistent with bystander recruitment of virus-specific CD8^+^ cells into the tumor bed after periodic stimulation due to reactivation of latent infections^31,62^. Finally, *S1PR5*^High^ clones were broadly distributed across tumor and adjacent normal tissue with no clear enrichment for the assayed antigen classes (Fig. 4h), suggesting an additional axis of infiltrating CD8^+^ T cells that should be explored further.

### Alignment of clonal embeddings shows conserved CD8^+^ T cell lineages across cancer types

In addition to analysis of clonal variation in individual datasets, it is often important to identify patterns of clonal variation recur across biological contexts. Here we asked if the patterns of clonal variation observed within the CD8^+^ T cells identified in NSCLC generalize to other tumor types. To do so we first used clone2vec to analyze datasets from NSCLC by Liu et al.^66^, colorectal cancer by Chen *et al.*^60^, head and neck squamous cell carcinoma (HNSCC) by McCord *et al*.^67^, as well as glioblastoma and a panel of brain metastases of different origins by Wang *et al*.^68^ Across these diverse datasets, the clonal embedding space frequently exhibited a similar organization, with clonal clusters marked by overlapping gene expression signatures (Fig. 5a-c and Supplementary Fig. 12), suggesting that major CD8^+^ clonal programs are not unique to NSCLC.

**Figure 5.**
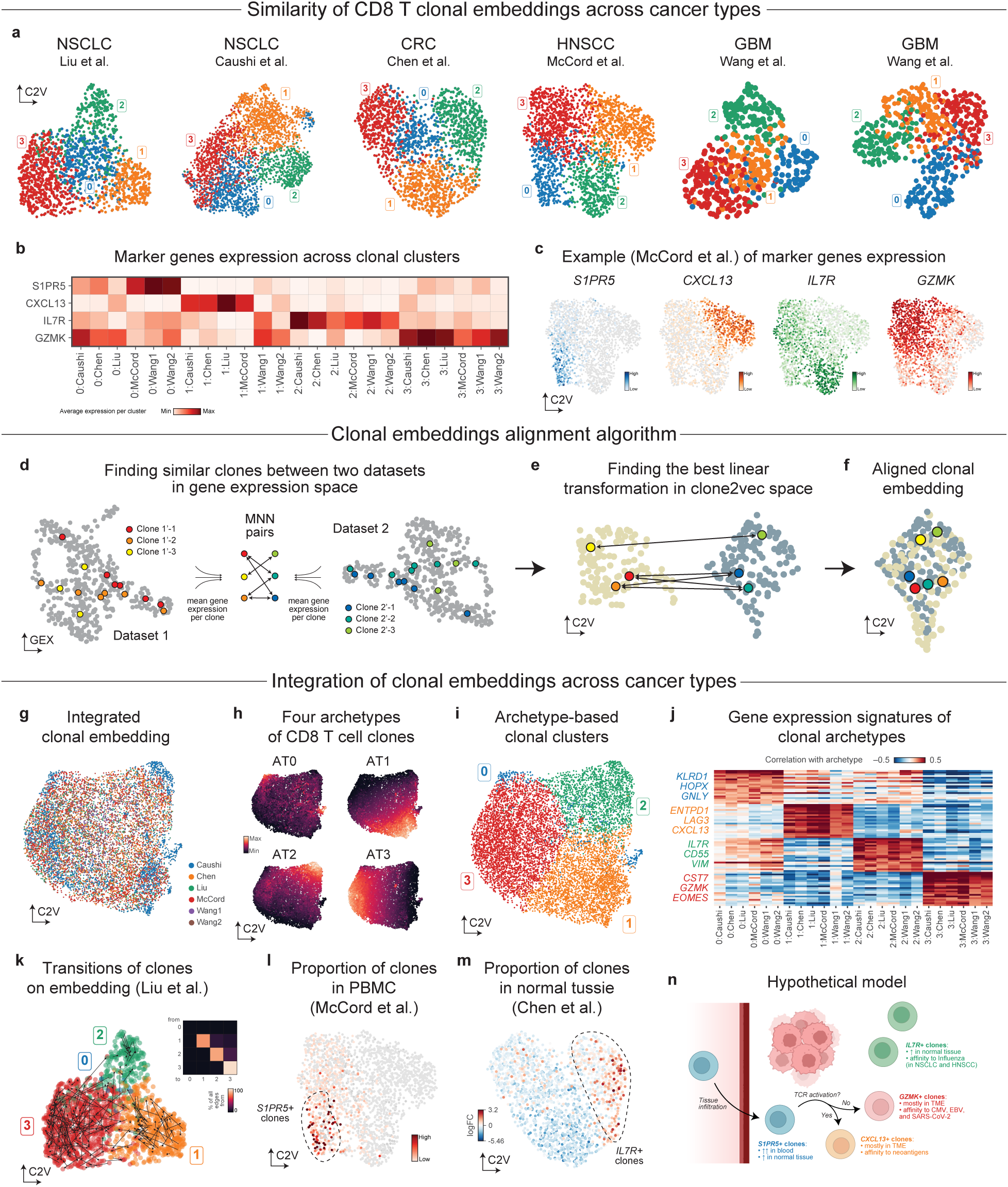
Joint analysis of clonal embeddings reveals recurrence of archetypal CD8^+^ T cell lineages across cancer types. **a**. clone2vec UMAPs of adaptive CD8^+^ T cell clones from the microenvironments of multiple cancers (left to right: lung cancer, Liu et al.; lung cancer, Caushi et al.; colorectal cancer, Chen et al.; head and neck cancer, McCord et al.; glioblastoma, Wang et al., CITE-Seq cohort; glioblastoma, Wang et al., scRNA-seq cohort), colored by archetype-based clusters; cluster numbers were manually matched across datasets based on marker gene expression. **b**. Heatmap of mean marker gene expression in clones from each archetype-based cluster across datasets. **c**. clone2vec UMAPs of CD8^+^ T cell clones from head and neck cancer (McCord et al.), colored by expression of *S1PR5* (archetype 0), *CXCL13* (archetype 1), *IL7R* (archetype 2), and *GZMK* (archetype 3). **d-f**. Schematic of the clonal embedding integration algorithm: (**d**) MNN identification between pairs of datasets using clone-averaged gene expression profiles; (**e**) hierarchical pairwise integration in clone2vec space using the previously identified MNNs; (**f**) the resulting integrated clonal embedding. **g**. Integrated clone2vec UMAP from the integration of the embeddings in (a), colored by cohort of origin. **h**. Integrated clone2vec UMAP arranged as a 2×2 grid (top row: archetypes 0, 1; bottom row: archetypes 2, 3), colored by integrated archetype weights. **i**. Same UMAP as (h), colored by archetype-based clusters. **j**. Heatmap of the minimum across-dataset Pearson correlation between per-clone average gene expression and each archetype’s weight. **k**. clone2vec UMAP of CD8^+^ T cell clones from lung cancer (Liu et al.), colored by integrated archetype-based clusters; arrows connect the same clone across timepoints (before and after anti-PD1 therapy). Inset (top right): heatmap of the relative proportion of transitions between clonal clusters across timepoints, normalized row-wise. **l**. clone2vec UMAP of CD8^+^ T cell clones from head and neck cancer (McCord et al.), colored by clonotype frequencies of matched clones in PBMCs. **m**. clone2vec UMAP of CD8^+^ T cell clones from colorectal cancer (Chen et al.), colored by logFC of clonotype frequencies between tumor and adjacent normal tissue. **n**. Schematic of a hypothetical model of T cell dynamics consistent with the observed archetypal lineages.

In principle, joint analysis of multiple datasets could proceed by integrating all single cells across studies and recomputing a unified clonal embedding. In practice, integrating T cells across diverse tumor types is challenging and risks overcorrecting lineage-specific differences. We therefore introduced a method to align clonal spaces directly, taking advantage of their algebraic structure (see Methods). Briefly, the approach used mutual nearest neighbors^69^ to match similar clones across datasets based on their average gene expression profile (Fig. 5d), and then found the best linear transformation to optimize their alignment in the joint latent space (Fig. 5e-f). Such a procedure guarantees that the algebraic structure of the individual clonal spaces will hold even after an integration.

The resulting integration of six different clonal embeddings has recapitulated the structure of the embedding initially observed for the NSCLC cohort Caushi *et al.* (Fig. 5g), described by four major archetypes associated with similar transcriptional signatures (Fig. 5h-j). Similarly, the patterns of clonal variation observed in analysis of other individual datasets were also well captured by the archetypes identified in the joint analysis (Fig. 5j and Supplementary Fig. 13). The robust recurrence of clonal variation patterns across diverse cancer types and tissues indicates that these archetypes capture general behavior of CD8^+^ T cells rather than specifics of tumor microenvironment in different tissues.

When aligning diverse datasets, it is also possible to relax the analytical constraints on clonal space, which can yield more structured clonal embeddings. Some of the existing methods allow decoupling the space used to establish similarity between datasets and the space where the integration is actually performed. For example, applying Seurat v5^70^ to integrate multi-cancer CD8+ T cell lineage data, shows an embedding that is consistent with the archetypes described earlier, but has a more pronounced cluster structure (Supplementary Fig. 14). An alignment preserving linear constraints, however, is more appropriate for the archetypal analysis.

Next, we used complementary strengths of different datasets to further characterize the clonal archetypes and improve the interpretation of these patterns. Liu *et al.* NSCLC study profiled site-matched tumors before and during anti-PD1 combination therapy, enabling temporal tracking of the same clonal lineages across treatment^66^. Examining whether treatment induces systematic transitions between archetypes, we found that most clones stayed in the same archetype (Fig. 5k). This illustrates that while anti-PD-1 therapy can dramatically alter clonal frequencies, shifts in the phenotypic properties of the clones are rare. The HNSCC data from McCord *et al.* included bulk TCR sequencing from matching PBMC samples. We found that *S1PR5*^High^ clones were significantly enriched in blood (Fig. 5l). In the colorectal cancer by Chen *et al*. we evaluated the extent of clonal sharing between tumor and adjacent normal tissue, we found enrichment of the *IL7R*^High^ clones, recapitulating the trend observed in the Cauchi *et al.* NSCLC dataset (Fig. 5m, Fig. 4g).

In summary, we demonstrate that a clone-centric perspective, enabled by clone2vec, provides a robust system-level view of the tissue that is not readily accessible from cell-level analyses alone. Furthermore, by aligning clonal embeddings across cohorts while preserving their algebraic structure, we enable integrative analysis that uses the complementary strengths of individual datasets to make generalizable conclusions about patterns of cell behavior.

## Discussion

As lineage-coupled single-cell profiling becomes more common, effective analysis will likely involve multiple computational strategies. Clone2vec implements one such strategy, centered on constructing a continuous clonal embedding to enable an unbiased, top-level exploration of prevalent clonal variation. The approach is focused on clones as indivisible analysis units (Supp. Fig. 14). By projecting clones into an informative low-dimensional space, clone2vec allows investigators to identify distinct clonal behaviors and follow them up by characterizing specific fate distributions and associated molecular programs. These embeddings are sensitive not only to which cell types are present within a clone, but also to their relative proportions and detailed states, allowing clone2vec to capture finer aspects of clonal variation than approaches based solely on discrete fate membership. This makes it possible to detect subtle shifts in lineage balance, partial biases, and graded differences between clones that would otherwise be overlooked. The resulting embedding provides a high-resolution description of clonal identity that can be linked directly to specific gene expression features, positional and anatomical coordinates, or developmental timing. In this way, clonal variation can be analyzed not as a set of coarse, predefined classes, but as a continuous and biologically interpretable landscape spanning lineage output, molecular state, spatial context, and temporal progression.

Fundamentally, comparing two clones amounts to comparing their distributions over the cellular expression manifold. Although many methods exist for comparing distributions based on samples, lineage-traced scRNA-seq data pose distinct challenges, stemming from small clone sizes, uneven clone-size distributions, and stochastic noise. Under these sparse regimes, the skip-gram approach implemented by clone2vec is significantly more robust than more common metrics like optimal transport or maximum mean discrepancy, allowing for a more stable recovery of the underlying biological structure.

An important illustration of the robustness of the clone2vec approach is the pan-cancer recurrence of the same four CD8^+^ T cell clonal subpopulations across a diverse range of solid tumors (Supp. Fig. 12-13); in the absence of ground-truth labels for clonal architecture, this cross-cohort reproducibility across independently generated datasets also serves as a real-data benchmark for the method. It suggests a highly conserved, multi-compartment model of TME infiltration and clonal fate determination. We interpret the *S1PR5*^High^ archetype as the circulating precursor or recently infiltrated pool, consistent with its enrichment in blood and the experimental evidence that *S1PR5* limits durable tissue residency^71^. Following tissue infiltration, clonal trajectories appear to bifurcate based on local TCR engagement. Clones recognizing tumor neoantigens are driven into the *CXCL13*^High^ archetype, a state characterized by chronic activation and canonical exhaustion markers (*e.g. ENTPD1*, *CD39*, *CTLA4*). Conversely, infiltrating clones lacking tumor-antigen affinity assume distinct bystander roles likely maintained by local inflammatory cytokines rather than direct tumor-antigen stimulation. The *IL7R*^High^ archetype, enriched in adjacent normal tissue and associated with influenza-reactive clonotypes, is most consistent with a tissue-associated memory-like bystander population^62,72^. The *GZMK*^High^ archetype may instead reflect a more inflammatory intratumoral bystander state, mobilized by cytokine-rich conditions within TME. While the enrichment for CMV- and EBV-reactive clonotypes suggests a potential contribution from latent antiviral memory, it may represent a broader *GZMK*^+^ CD8^+^ tumor-infiltrating program previously observed across multiple cancers^62,68,73^. This interpretation is also consistent with the observed scarcity of transitions between archetypes during anti-PD-1 therapy, as one would not expect checkpoint blockade to alter archetype membership if it is fundamentally determined by inherent TCR affinities.

The clone2vec approach should generalize well to other modalities and tracing techniques. The presented examples are mostly focused on the available scRNA-seq datasets, however, the graph-based approach employed for assessing molecular similarity between cells is directly applicable to analysis of lineage-tracing coupled to other modalities (*e.g.* ATAC-seq, *etc.*). In terms of tracing techniques, the examples are focused on applications to static barcodes, including multiple transgenic barcode designs, as well as endogenous TCR “barcodes”. Expanding the approach to dynamic barcoding techniques, both prospective^8,74^, as well as retrospective^75,76^, would be an exciting future direction.

The biological phenomena captured by any clonal embedding depend on the experimental design, particularly on the timing of barcode induction. In developmental studies, for example, barcodes can be introduced early, before substantial commitment has occurred. The clonal embeddings will then capture variation in progenitor competence and differentiation bias^8,38^. In contrast, TCRs are formed together with cell type commitment, and the clonal variation will reflect plasticity and clonal restrictions between different states of a single, committed cell type. In this setting, clone2vec is advantageous because it does not depend on predefined cell annotations, but instead quantifies clonal similarity directly from cell distributions in continuous expression space, allowing unbiased exploration of state structure and enabling potential discovery of cell subpopulations distinguished by their clonal relationships.

A key strategy to follow up on the observed clonal variation patterns is to link them to the underlying molecular signatures. We have described several such common test scenarios. For example, in a differentiating context, one would commonly test for genes in progenitor populations associated with downstream lineage biases (Fig 2g,h). Conversely, testing for expression associations in a distinct mature cell population can be used to reveal elaborate coordination of cell behaviors in a tissue, such as the illustrated difference in the state of monocyte cells in clones that do or do not contribute to neutrophils (Supplementary Fig. 8a-h)^19^. Finally, clone2vec also implements global association tests, to identify gene expression patterns in all or select cell subpopulations that explain as much of the overall observed clonal variation as possible (Fig. 2m-o). When the goal is to link early progenitor state to later fate, dedicated frameworks such as CoSpar^28^ may be preferable because they explicitly exploit manifold continuity and can borrow information from unlabeled cells, thereby increasing the effective sample size for fate-bias inference. Clone2vec should therefore be viewed as complementary to these approaches: it is strongest as an annotation-independent, exploratory representation of clonal variation that supports hypothesis generation, cross-context comparison and subsequent targeted testing.As lineage-enabled assays continue to scale, we hope clone2vec will be instrumental in extracting biological insights from such data.

## Data availability

No new data were generated or analysed in this study. Previously published datasets used in this work are available from the following sources: Haan et al., ArrayExpress (E-MTAB-14817); Ireland et al., Zenodo (10.5281/zenodo.15857303); Weinreb et al., via the CoSpar package (cospar.datasets.hematopoiesis_130K); Sureshchandra et al., Zenodo (10.5281/zenodo.18868813 and 10.5281/zenodo.13119615); Luoma et al., Gene Expression Omnibus (GSE144469); Chen et al., GEO (GSE236581); Caushi et al., GEO (GSE176022); Liu et al., GEO (GSE179994); McCord et al., GEO (GSE287301 and GSE300147); and Wang et al., Zenodo (10.5281/zenodo.10672442).

## Code availability

The clone2vec Python package is available on GitHub (https://github.com/kharchenkolab/clone2vec) together with the code for the reproducibility of the analysis (https://github.com/kharchenkolab/clone2vec_analysis).

## Acknowledgements

We thank Emma Andersson (Karolinska Institutet) for helpful discussions during manuscript preparation, and her integral role in a companion manuscript that inspired this work. I.A. and S.I. were supported by the ERC Synergy grant (KILL-OR-DIFFERENTIATE), Swedish Research Council, Paradifference Foundation, Bertil Hallsten Research Foundation, Cancer Foundation in Sweden, Knut and Alice Wallenberg Foundation, Austrian Science Fund. A.G.E. was supported by the National Institute of Dental and Craniofacial Research of the National Institute of Health under award number 1F32DE029662, and the Swedish Research Council Starting Grant in Medicine and Health (#2024-03554). P.V.K. was supported by the ERC Synergy grant (KILL-OR-DIFFERENTIATE) and the Visiting Fellowship from the Austrian Academy of Sciences.

## Contributions

S.I. performed data analysis for the manuscript. A.G.E. helped with biological interpretation of the developmental data. I.A. secured funding and conceptually supervised the study. P.V.K. technically and conceptually supervised the study. P.V.K., S.I., and I.A. drafted the first version of the manuscript. All authors jointly conceived the main idea of this study and participated in discussing and reviewing the manuscript.

## Methods

### clone2vec algorithm

At its core, clone2vec can be viewed as an algorithm for the multinomial matrix factorization of a clone-by-clone nearest-neighbor co-occurrence matrix. In more detail, the algorithm consists of the following steps (for more formal description see Supplementary Note).

#### 1. Construction of an appropriate gene expression embedding

The algorithm relies on two inputs: (a) a gene expression embedding and (b) clonal labels. The gene expression embedding is a low-dimensional representation of cells (typically 15-30 dimensions) that captures the biologically relevant variance in the data. In standard pipelines, it can be PCA computed from gene expression; for multi-batch datasets, it can be a batch-corrected embedding such as Harmony^77^, an MNN-corrected PCA space (as in Seurat^70^), or a VAE latent representation (as in scVI^78^). Proper batch correction in this space is essential. Otherwise, cells from the same batch will group together in the final clonal embedding because of residual batch effects rather than shared biology – the batch signal propagates through the kNN graph into the clone-level representation. The clonal labels form a single vector assigning a clonal identity to each cell. For evolving-barcode lineage tracing, an appropriate level of the lineage tree must be chosen to collapse cells into a single flat set of clones.

#### 2. kNN graph construction and clonal neighborhood aggregation

Using the batch-corrected gene expression embedding, we build a k-nearest-neighbor (kNN) graph on clonally labeled cells (for example, with *k* = 15). From this graph, for each cell we count how many of its k neighbors belong to each clone, producing a vector of length *C*, where *C* is the number of unique clones in the dataset. We then aggregate these per-cell vectors to the clone level by summing them across all cells of the same clone. The result is a *C × C* matrix whose entry *(i, j)* is the total number of neighbors from clone *j* observed in the gene expression neighborhoods of all cells from clone *i*.

#### 3. Skip-Gram

This step can be described in two mathematically equivalent ways^36^ – via a Skip-Gram neural network or via multinomial GLM-PCA – and for now we focus on the Skip-Gram formulation. Skip-Gram is a simple neural network with a single hidden layer of size *z* (the number of latent dimensions in clone2vec; typically *z* is 10) and input and output layers of size *C*. The activations in the hidden and output layers are linear. With no further modifications and an MSE loss, this architecture is equivalent to PCA^79^. Non-linearity is introduced by applying a softmax to the output layer, which transforms the real-valued output scores into a probability distribution over clones. These predicted probabilities are compared with the empirical probabilities of observing one clone in the gene expression neighborhood of another, and the model is trained by minimizing the resulting negative log-likelihood. Training is performed in shuffled mini-batches, using individual pairs of clones as elementary training examples; this is what makes the present implementation closer to a Skip-Gram network than to GLM-PCA, where the full matrix is typically fit at once. After convergence, the input-to-hidden weight matrix is used as the latent representation of clones.

### Poisson GLM-PCA as an alternative to Skip-Gram

As noted above, Skip-Gram can be equivalently formulated as multinomial GLM-PCA (Generalized Linear Model PCA) – the same model is also referred to as multinomial EPCA (Exponential-family PCA) in the earlier literature^37^. In the multinomial formulation, efficient parallel optimization is limited because the softmax normalization couples all output units. In natural language processing, negative sampling – a procedure in which the full softmax is replaced by a binary classification against a small set of sampled negative examples – is routinely used for large corpora; its computational advantages are most pronounced when the output vocabulary is very large and the embedding dimension is also large (e.g., 200-300), and are minimal or even detrimental at the small embedding dimensions used in clone2vec (*z = 10*).

If the multinomial likelihood is replaced by a Poisson likelihood, each output unit becomes conditionally independent of the others, which enables efficient parallel optimization via Alternative Poisson regression, as it’s shown in fastglmpca^46^ (now implemented in fastglmpca-py). With column-wise and row-wise normalization, the resulting embeddings become comparable to those of the multinomial model, making the Poisson variant a fast alternative to standard clone2vec for datasets whose size would otherwise be prohibitive. In ordinary cases, however, we recommend the vanilla multinomial clone2vec, as it more faithfully captures the compositional structure of the neighborhood counts and offers stronger theoretical guarantees on the resulting solution.

### Clustering and archetypal analysis in clone2vec space

The output of clone2vec – a latent representation of each clone – can serve as input to the standard downstream analyses developed for single-cell RNA-seq data. Throughout the paper, we use UMAP^80^ visualizations of the clone2vec space to inspect its local structure and, in some cases, trajectory inference methods to characterize continuous axes of variation in clonal composition. For discrete grouping, we recommend community-detection algorithms such as Leiden^81^.

When the boundaries between clonal clusters are not well defined – a situation that arises frequently in practice, due to the presence of small clones, genuinely fuzzy boundaries between cell types, and the inherently continuous nature of compositional variation across clones – archetypal analysis provides a complementary view of the heterogeneity in clonal behavior. In brief, archetypal analysis fits a convex polytope with *A* vertices, called archetypes, to the data under two requirements: each data point is approximated as a convex combination of the archetypes, and each archetype is itself a convex combination of data points^82^. The archetypes therefore lie near the extremes of the data distribution and act as interpretable reference points, with every clone expressed as a mixture of them. Because the clone2vec space preserves the compositional semantics of clones in its geometric structure, the resulting archetypes tend to recover stereotypical patterns of clonal behavior that jointly account for the bulk of the observed variation. After this analysis, each clone is represented by its coordinates in archetype space – by the weights of its convex combination over the A archetypes. These coordinates can be reduced to a hard, cluster-like assignment by labelling each clone with the archetype that carries the largest weight.

### Cluster-level gene expression associations

Once the clonal variation of interest has been characterized – by building a trajectory and assigning a pseudotime value to each clone, by clustering, or by archetypal analysis – the resulting groupings or orderings can be explored with classical differential expression techniques. We recommend treating clones, rather than individual cells, as the units of replication in these tests: for each clone, gene expression is averaged across the cells of the cell type of interest, and differential expression is then performed on the resulting clone-level pseudobulk profiles.

### Cluster-free gene expression associations analysis

To exploit the continuous and multidimensional nature of clone2vec embeddings, rather than collapsing them into discrete labels, we provide a framework for cluster-free analysis of the association between gene expression and position in clone2vec space, based on supervised PCA^42^. The idea is to use clone2vec coordinates to define which clones are close to each other, and then to look for patterns of gene expression that vary smoothly across that neighborhood structure – i.e. patterns in which clones that are close in clone2vec space also tend to have similar expression.

Concretely, we build a k-nearest-neighbor graph on the clones in clone2vec space, and then identify the directions in gene expression space along which neighboring clones are most similar. These directions form a small set of graph-aware principal components, each corresponding to a coordinated program of genes whose expression changes gradually as one moves across the clone2vec manifold. Each component comes with an associated measure of how much of the smooth, neighbor-structured variation it captures, which allows the components to be ranked by relevance and truncated as in ordinary PCA. Top-ranked components correspond to the broadest patterns of variation – smooth gradients that span large regions of the clone2vec space and reflect gene programs differentiating major clonal behaviors – while lower-ranked components capture progressively finer-scale, more localized patterns, down to subtle expression differences among nearby clones.

As a univariate complement to this joint decomposition, we compute Moran’s I^83^ for each gene – a standard measure of the extent to which a signal is organized by an underlying graph structure – and test, for every gene, whether its expression is more spatially structured in clone2vec space than would be expected by chance. The resulting per-gene p-values are adjusted for multiple testing with the Benjamini-Hochberg procedure. Together, the graph-aware principal components describe the dominant coordinated patterns of expression variation across the clone2vec manifold, while Moran’s I flags individual genes whose expression is significantly shaped by clone2vec geometry.

### Horizontal alignment of clonal embeddings

Jointly analyzing several clone2vec embeddings from different cohorts is an additional computational challenge that arose during method development. Standard integration of multiple single-cell datasets at the cell level can erase biologically meaningful variation and complicate interpretation of the resulting embedding. To avoid this, we developed an integration procedure that operates directly on clonal embeddings.

The central difficulty is that clone2vec is trained independently for each cohort, and its latent space is identifiable only up to an arbitrary linear transformation: the axes carry no intrinsic meaning in terms of cell-type composition, and coordinates are not directly comparable across cohorts. An external reference is therefore required to match clones between datasets. We use the average gene expression per clone together with a mutual nearest neighbor (MNN) strategy^69^, through the following multi-step procedure:

1. Per-clone average gene expression is computed by averaging the log-normalized expression of all genes across the cells assigned to each clone.
2. Clones from all cohorts are concatenated into a single dataset in which each clone is treated as a pseudo-cell.
3. Highly variable genes are identified in a batch-aware manner, with cohort as the batch variable.
4. Expression is scaled per cohort prior to PCA.
5. PCA is computed on the scaled expression matrix.
6. The resulting PCA space is integrated across cohorts with Harmony.
7. Mutual nearest neighbors are identified between every pair of cohorts in the Harmony-corrected space.

The matched clone pairs – which can be referred to as anchors, following Seurat’s terminology^84^ – are then used to bring the clone2vec coordinates of different cohorts into a common frame. Alignment is performed hierarchically: cohorts, or groups of already-aligned cohorts, are merged in order of decreasing number of shared anchors, so that the most strongly connected cohorts are integrated first. At each merge step, the smaller cohort (or group) is mapped onto the larger one by a weighted affine transformation – a linear map followed by a translation – whose parameters are obtained by weighted least squares on the anchor pairs, using the weights defined above. The final result is a single aligned clone2vec embedding in which clones from all cohorts share a common coordinate system.

### Details of benchmarking

To benchmark the ability of clone2vec to capture continuous changes in clonal composition, we generated a 2D “gene expression” space with two well-separated clusters (A and B). After that, we generated clones using cells from this gene expression space so each clone consisted of 10 cells and had exactly *n* (*n* = 0, 1, …, 10) cells from cluster A and *10 − n* cells from cluster B; for each n the procedure was repeated 500 times, resulting in 5500 clones in total. Clonal embedding was built with only two latent dimensions, and the first principal component of the clone2vec space was used to find a correlation between the position of the clone on a clonal embedding and the proportion of cell type A in the clone. In the case of Poisson GLM-PCA benchmarking, the first GLM-PC was used to estimate the correlation.

To evaluate the ability to capture visibly distinct clonal behaviors and support semantic geometry, we generated a 2D “gene expression” space with three clusters (A, B, and C); after that, we generated clones (each of size 15) consisting of three “portions” (of 5 cells each): each “portion” can be from cluster A, B, or C. Depending on which clusters were involved in the clone construction, we labeled these clones as “AAA” (for clones with 15 cells from cluster A), “ABB” (for clones with 5 cells from A and 10 cells from B), “ABC” (for clones with 5 cells from each cluster), and so on. Clonal embedding with a latent space of size 2 was built based on 500 clones of each type (totaling 5000 clones), and the resulting geometry was evaluated visually.

For more complex clonal distributions (“Rings and Crosses” benchmark), we constructed a 2D “gene expression” space containing four concentric rings and six crosses rotated at different angles (0°, 15°, 30°, 45°, 60°, and 75°). Clones of size 30 were generated by sampling cells from these specific geometric structures (one clone is sampled from one specific structure). For each of the ten structures (four rings and six crosses), 300 clones were simulated, resulting in 3000 clones in total. After that, the dataset was subsampled into subsets consisting of 1%, 5%, 10%, 25%, and 50% of the original data. Using these subsets, clonal embeddings for clones with size ≥ 2 were built. Using the same set of clones and the location of cells from them in the original 2D “expression space”, we calculated a variety of different metrics (such as MMD, Sinkhorn divergence, Energy distance, and cluster-based composition). Шт situations where the method returns distances (MMD, Sinkhorn, Energy) instead of vector representations, we used these distances directly for kNN graph construction and clustering. The performance was evaluated by performing Leiden clustering on the clonal embedding and comparing the results to the ground truth labels using the Adjusted Rand Index (ARI), searching for the best-matching clustering resolution starting from 0.1 and ending with 2 with a step of 0.1.

### Details of the analysis of each individual dataset

Unless stated otherwise, processing started from raw counts, which were library-size normalized (such that the total count per cell summed to 10,000 UMIs), after which 3,000 highly variable genes were selected in a batch-aware manner. These genes were then scaled and used for PCA (typically reduced to 20 principal components). When multiple batches were present, Harmony77 was used for batch correction. The Harmony-corrected space was used to build a kNN graph between clonally labelled cells (from clones of size ≥ 3, k = 15) and to calculate the nearest neighborhood for each clone. This clonal neighborhood matrix was then fed into a Skip-Gram clone2vec model with 10 latent dimensions. The resulting clone2vec space was used to build a clone-to-clone similarity kNN graph (k = 15) for subsequent clustering and dimensionality reduction with UMAP. For differential expression analysis, each clone was represented by the average expression across all cells of the cell type of interest, and a Welch’s t-test with Benjamini-Hochberg multiple testing correction was applied to identify genes whose expression differed significantly between groups of clones. When a subset of clones was selected (e.g., CD8 T cells from all T cells), this was typically done after the main embedding, which contained all cells of interest, had been built. Assuming that 10 clone2vec dimensions are sufficient to capture even fine differences between clones, we used only the clones of interest to build a kNN graph on this subset for further clustering and dimensionality reduction.

T cell clonotypes were identified from AIRR/CellRanger VDJ-like files, using a full match of both chains’ CDR3 nucleotide sequences as the criterion for grouping cells into clones. Clones containing more than one TRB sequence were excluded; in the case of multiple TRA sequences per cell, all TRA sequences were required to match within the clone. All procedures were performed using the scirpy^85^ Python package. Antigen lookup in VDJdb^65^ was also performed with scirpy, using the amino acid sequences of the CDR3 regions of both alpha and beta chains. Archetypal analysis was performed using the ParTIpy^86^ Python package, which implements the PCHA^87^ algorithm. Four archetypes were selected based on explained variance, information criteria, and bootstrap stability. In cases where datasets consisted of multiple biopsies per patient (e.g., tumor / LN / metastasis), only T cells from the TME were used to build the clonal embedding.

**Haan et al.**^38^ Data were obtained from ArrayExpress (E-MTAB-14817). Neurons were subselected from the whole dataset based on the expression of *Tubb3* and *Sox1*. Clone-type annotation was performed based on the average expression across all cells of each clone. In the cluster-free analysis, each clone was represented by multiple data points corresponding to the number of distinct cell types present in that clone (e.g., a clone containing 3 cell types was represented by 3 data points sharing the same clone2vec position). In addition, the mean expression per cell type, computed across all clones, was subtracted from the cell-type-specific expression vector of each individual clone. This step, together with the previous one, was performed to remove the influence of cell type markers on the SPCA procedure. The final SPCA was then built using clone2vec-based similarity between the resulting data points (including multiple replicates of the same clone originating from different cell types) and the scaled expression of each cell-type-restricted clone. Trajectory inference on the clone2vec embedding was performed with the ElPiGraph^88^ algorithm implemented in scFates^89^, using 15 nodes for principal curve fitting based on all 10 dimensions of the clone2vec latent representation.

**Ireland et al.**^40^ Data were obtained from Zenodo (10.5281/zenodo.15857303). The scVI latent space from the original object (“X_scVI_1.2”) was used to build a kNN graph, which was subsequently used for clonal embedding construction. Leiden clustering with resolution 0.75 was performed to obtain clonal clusters. Gene signature scores were calculated with the score_genes scanpy function, using *Epcam*, *Krt8*, and *Vwf* for the epithelial score (E-score); *Stmn2*, *Nrxn3*, *Dcx*, *Zic5*, *Dmd*, *Negr1*, and *Tenm2* for the neuronal progenitor score (Nprog-score); and *Stmn2*, *Nrxn3*, *Dcx*, *Tshz2*, *Myt1l*, *Ryr3*, and *Plxna4* for the migratory neuron score (Nmig-score).

**Weinreb et al.**^19^ Data were obtained via the CoSpar Python package (cospar.datasets.hematopoiesis_130K). The PCA space from the original object was batch-corrected with Harmony. This Harmony-corrected space was used for all computations, while the original SPRING layout was used to visualize the distribution of cells in gene expression space. *Cd34*^+^ progenitors were selected based on smoothed expression of *Cd34* (kNN smoothing, 20 steps) with a threshold of 0.5 (a similar procedure was used in the original paper). Communities were identified with the Leiden algorithm at resolution 2. For the cluster-free analysis within the monocyte fate, all monocytes were masked, and neighbors of clones within this group were collapsed into a single “monocyte” entity (instead of counting individual clonal labels of the neighbors). This ensured that clonal heterogeneity within the monocyte fate itself did not influence the resulting clonal embedding (preventing SPCA from capturing heterogeneity within monocytes that is unrelated to clonal heterogeneity). For the ClonoCluster illustration, the algorithm was reimplemented in Python using the default parameters from the original paper. The final ClonoClusters were obtained by applying the Leiden algorithm to the ClonoCluster graph at resolution 1.

**Sureshchandra et al.**^45^ Data were obtained from Zenodo (10.5281/zenodo.18868813 and 10.5281/zenodo.13119615). The analysis mostly followed the procedure described earlier. T cell clonotypes were also obtained from the original data objects. A two-sided Mann-Whitney U test without multiple testing correction was used to estimate the significance of differences in fate proportions between PBMC parts within each clonal cluster.

**Luoma et al.**^61^ Data were obtained from GEO (GSE144469). The analysis mostly followed the procedure described earlier. Compositional analysis was performed using a two-sided Mann-Whitney U test without correction for multiple testing.

**Chen et al.**^60^ Data were obtained from GEO (GSE236581). The analysis mostly followed the procedure described earlier.

Common markers from the Luoma, Chen, and Sureshchandra datasets were identified by intersecting the upregulated and downregulated genes (|logFC| ≥ 1, two-sided Welch’s t-test with Benjamini-Hochberg correction, adjusted p-value < 0.05) from the differential expression analysis between tTreg-like and pTreg-like clones (using expression values only from the Treg portion of each clone).

**Caushi et al**.^31^ Data were obtained from GEO (GSE176022). The analysis mostly followed the procedure described earlier. For visualization, the authors’ provided UMAPs were used; for analysis, the latent representation of cells was rebuilt using the procedure described above.

**Liu et al.**^66^ Data were obtained from GEO (GSE179994). The analysis mostly followed the procedure described earlier. Clones from two timepoints (before and after anti-PD1 therapy) were treated independently.

**McCord et al.**^67^ Data were obtained from GEO (GSE287301 and GSE300147). The analysis mostly followed the procedure described earlier.

**Wang et al.**^68^ Data were obtained from Zenodo (10.5281/zenodo.10672442). In both cohorts, Seurat’s CCA integration in gene expression space was used to correct for batch effects. For the multimodal dataset, a WNN^41^ graph was built using the muon^90^ Python package. This WNN graph was then vectorized using Laplacian eigenvectors, and the resulting representation was fed into the clone2vec pipeline. The rest of the analysis mostly followed the procedure described earlier.

For further details, see the accompanying GitHub repository.

