## Supplementary Fig. for "Clonal embeddings allow exploratory analysis of lineage-resolved single-cell data"

**TABLE OF CONTENTS**


| | |
|---|---|
| Supplementary Figure 1 | Goals of clonal embeddings |
| Supplementary Figure 2 | Benchmark of clone2vec on continuous and discrete variation |
| Supplementary Figure 3 | Benchmark of clone2vec on complex clonal distributions ("rings and crosses") |
| Supplementary Figure 4 | Comparison of clone2vec to alternative methods for dropout stability |
| Supplementary Figure 5 | Details of the analysis of the Haan et al. dataset |
| Supplementary Figure 6 | Projection of individual clonal clusters (Leiden resolution = 2) onto the gene expression UMAP of CNS neurons from Haan et al. |
| Supplementary Figure 7 | Clonal analysis of the SCLC organoid differentiation dataset (Ireland et al.) |
| Supplementary Figure 8 | Clonal analysis of in vitro hematopoiesis from Weinreb et al. |
| Supplementary Figure 9 | Details of the analysis of PBMC and tonsil datasets from Sureshchandra et al. |
| Supplementary Figure 10 | Details of the analysis of colon T cell datasets from Chen et al. and Luoma et al. |
| Supplementary Figure 11 | Details of the analysis of T cells from NSCLC patients (Caushi et al.) |
| Supplementary Figure 12 | Per-archetype marker expression across cancer types |
| Supplementary Figure 13 | Integrated archetype weights across cancer types |
| Supplementary Figure 14 | Comparison of the proposed integration of clonal embeddings to a Seurat-based approach |
| Supplementary Figure 15 | Differences in purpose between clone2vec and ClonoCluster |

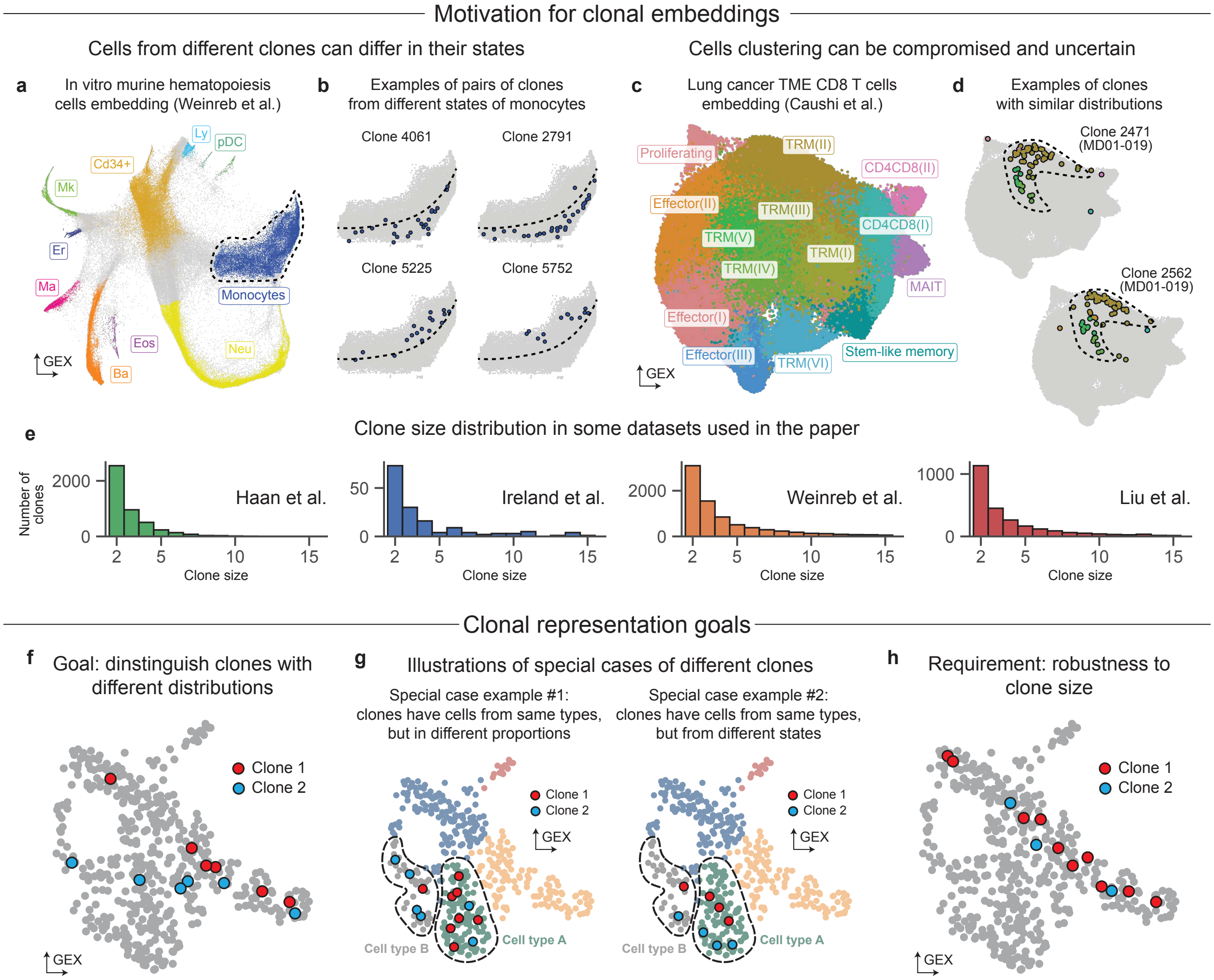


**Supplementary Figure 1**. *Goals of clonal embeddings*

**a**. Gene expression UMAP of in vitro murine hematopoietic cells from Weinreb et al., colored by cell types as defined in the original publication, with the dashed outline highlighting the monocyte compartment. **b**. Four example clones shown on the monocyte region of the UMAP from (a): clones 4061 and 2791 (top row) occupy a similar transcriptional substate, as do clones 5225 and 5752 (bottom row), yet the two pairs localize to distinct regions within the monocyte compartment. **c**. Gene expression UMAP of CD8 T cells from a lung cancer cohort (Caushi et al.), colored by the authors' cell types. **d**. Two example clones on the gene expression UMAP from (c), showing similar distribution patterns. **e**. Histograms of clone-size distributions in four clonally labeled datasets (left to right: Haan et al., Ireland et al., Weinreb et al., Liu et al.). **f-h**. Schematic gene expression UMAPs illustrating the main aims of the clonal representation task. (**f**) The primary goal of clonal embeddings is to construct a metric space for clones that is sensitive to differences in their distribution in gene expression space, including (**g**, left) compositional differences between clones and (**h**, right) subtle differences in expression space; the method must additionally be dropout-robust, as also illustrated in (**c**).

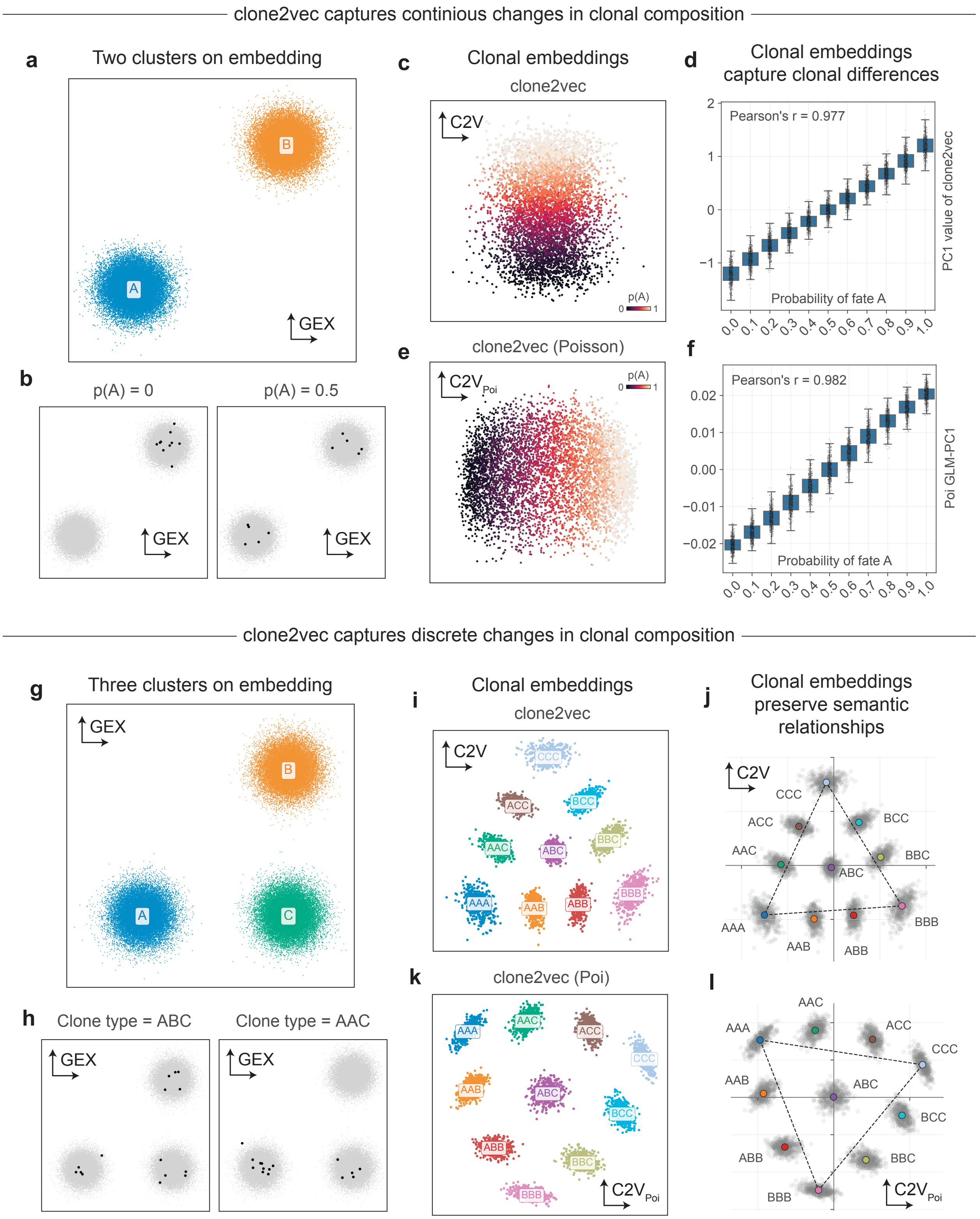


**Supplementary Figure 2**. *Benchmark of clone2vec on continuous and discrete variation*

**a**. Scatterplot of a simulated 2D gene expression embedding consisting of two clusters. **b**. Scatterplots of two example clones with different proportions of cells from cluster A (10 cells per clone). **c**. Resulting 2D clone2vec embedding from 5,500 clones, colored by the proportion of cells from gene expression cluster A. **d**. Box plot of the PC1 coordinate in clone2vec space, grouped by proportion of cluster A cells per clone. **e**. Resulting 2D Poisson clone2vec embedding from 5,500 clones, colored by the proportion of cells from cluster A. **f**. Box plot of the GLM-PC1 coordinate, grouped by proportion of cluster A cells per clone. **g**. Scatterplot of a simulated 2D gene expression embedding consisting of three clusters. **h**. Scatterplots of two example clones with different cluster compositions. **i**. Resulting 2D clone2vec embedding, colored by clone type. (Here and in (k), each letter in the clone-type name indicates 5 cells from the corresponding gene expression cluster.) **j**. Same embedding as (i), with dotted lines connecting the average coordinates of clone types "AAA", "BBB", and "CCC" to highlight the linear properties of the latent space. **k**. Resulting 2D Poisson clone2vec embedding, colored by clone type. **l**. Same embedding as (k), with dotted lines as in (j).

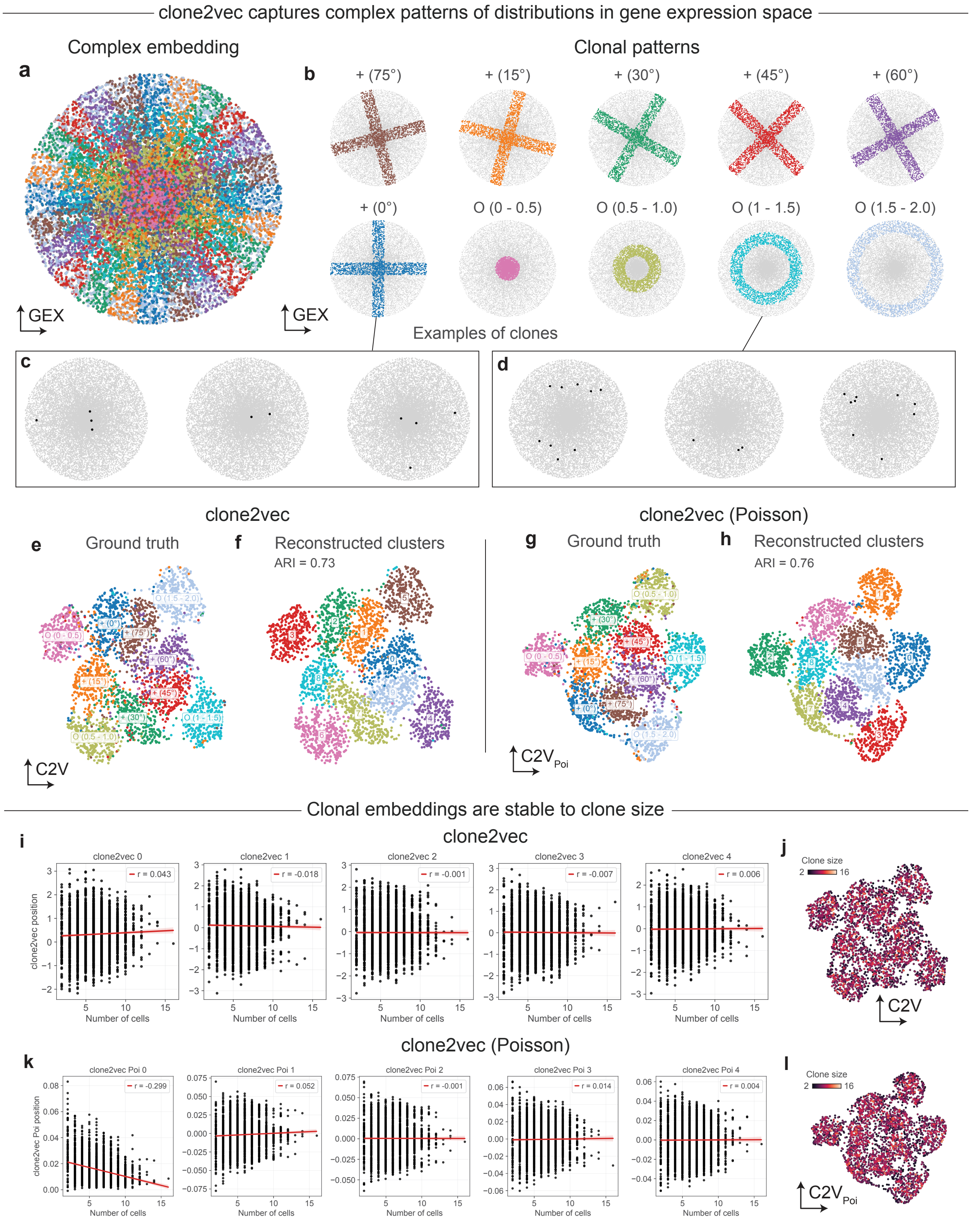


**Supplementary Figure 3**. *Benchmark of clone2vec on complex clonal distributions ("rings and crosses")*

**a**. Scatterplot of the simulated 2D gene expression space, colored by clone type. **b**. Illustration of each clone type used in the simulation. Initially, 300 clones of size 30 were generated for each clone type; the dataset was then subsampled to 20% of the original, yielding random clone sizes (range: 2-16). **c**. Examples of clones of type "+ (0°)" in gene expression space. **d**. Examples of clones of type "O (1-1.5)" in gene expression space. **e-f**. UMAP of the clone2vec representation, colored by (**e**) ground-truth clone type and (**f**) Leiden clusters at the best-matching resolution (ARI = 0.73). **g-h**. UMAP of the Poisson clone2vec representation, colored by (**g**) ground-truth clone type and (**h**) Leiden clusters at the best-matching resolution (ARI = 0.76). **i**. Scatterplots of clone2vec latent coordinates (y-axis) vs clone size (x-axis) with regression lines overlaid; correlations are negligible ($|r| < 0.05$) in all cases. **j**. Same UMAP as (e-f), colored by clone size. **k**. Scatterplots of Poisson clone2vec latent coordinates (y-axis) vs clone size (x-axis) with regression lines overlaid; correlations are negligible ($|r| < 0.05$) for all components except GLM-PC1 and GLM-PC2, with the largest observed for GLM-PC1 ($r \approx -0.3$). l. Same UMAP as (g-h), colored by clone size.

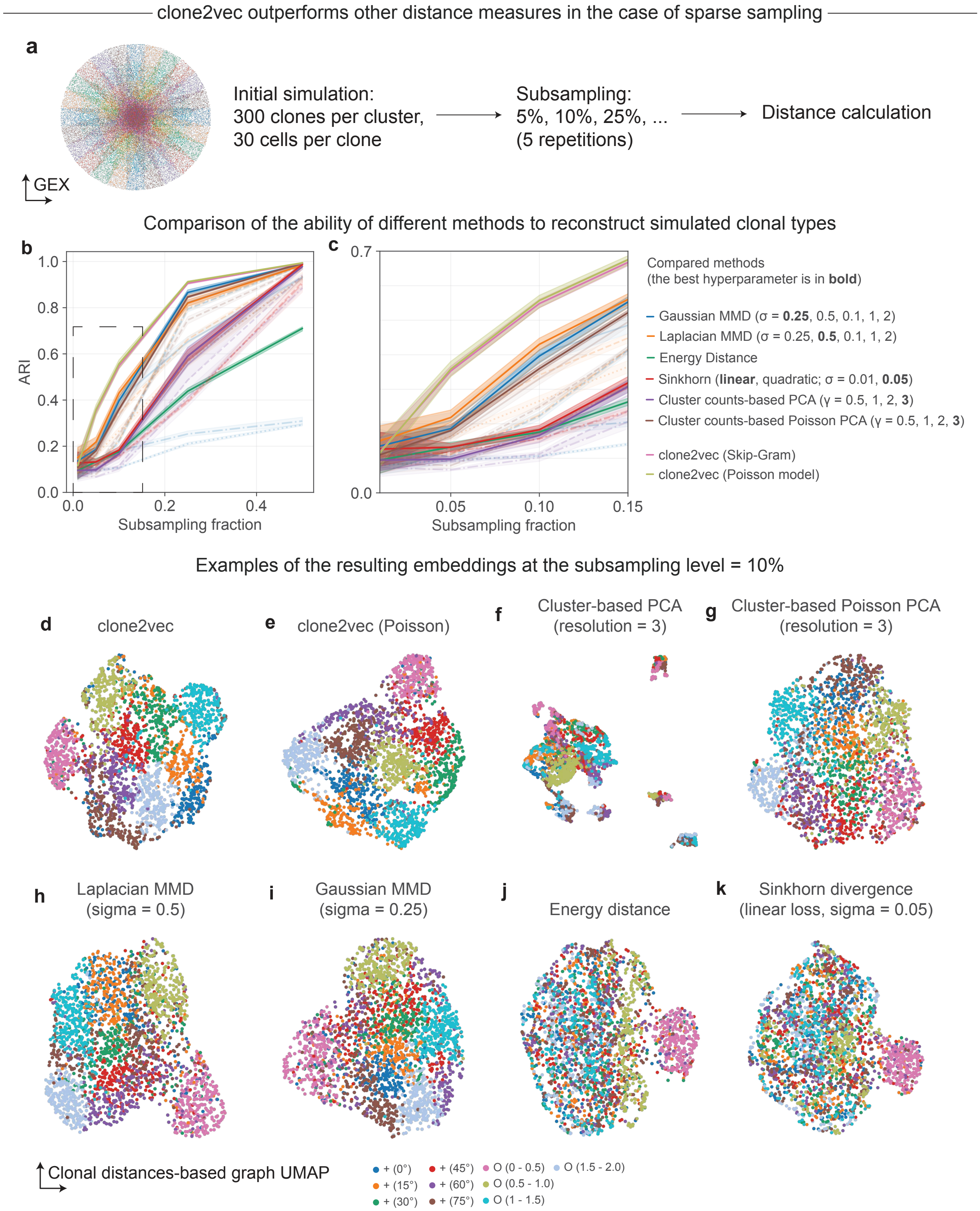


**Supplementary Figure 4**. *Comparison of clone2vec to alternative methods for dropout stability*
**a**. Schematic of the benchmarking procedure. Clones were generated as in Supplementary Figure 3 and subjected to varying levels of subsampling (subsampling fractions indicated on the figure), with 5 replicates per level. **b**. Line plot (lines, mean; shaded areas, 95% CI) of comparative performance across all methods, with the ARI of the best-matching

Leiden clustering resolution as the performance metric. **c**. Same as (b), zoomed in on small subsampling fractions. **d-k**. UMAPs based on vector representations (clone2vec (**d**), Poisson clone2vec (**e**), cluster-based PCA (**f**), cluster-based Poisson GLM-PCA (**g**)) or on kNN graphs constructed directly from inter-clone distances (Laplacian MMD (**h**), Gaussian MMD (**i**), Energy distance (**j**), Sinkhorn divergence (**k**)), colored by ground-truth labels.

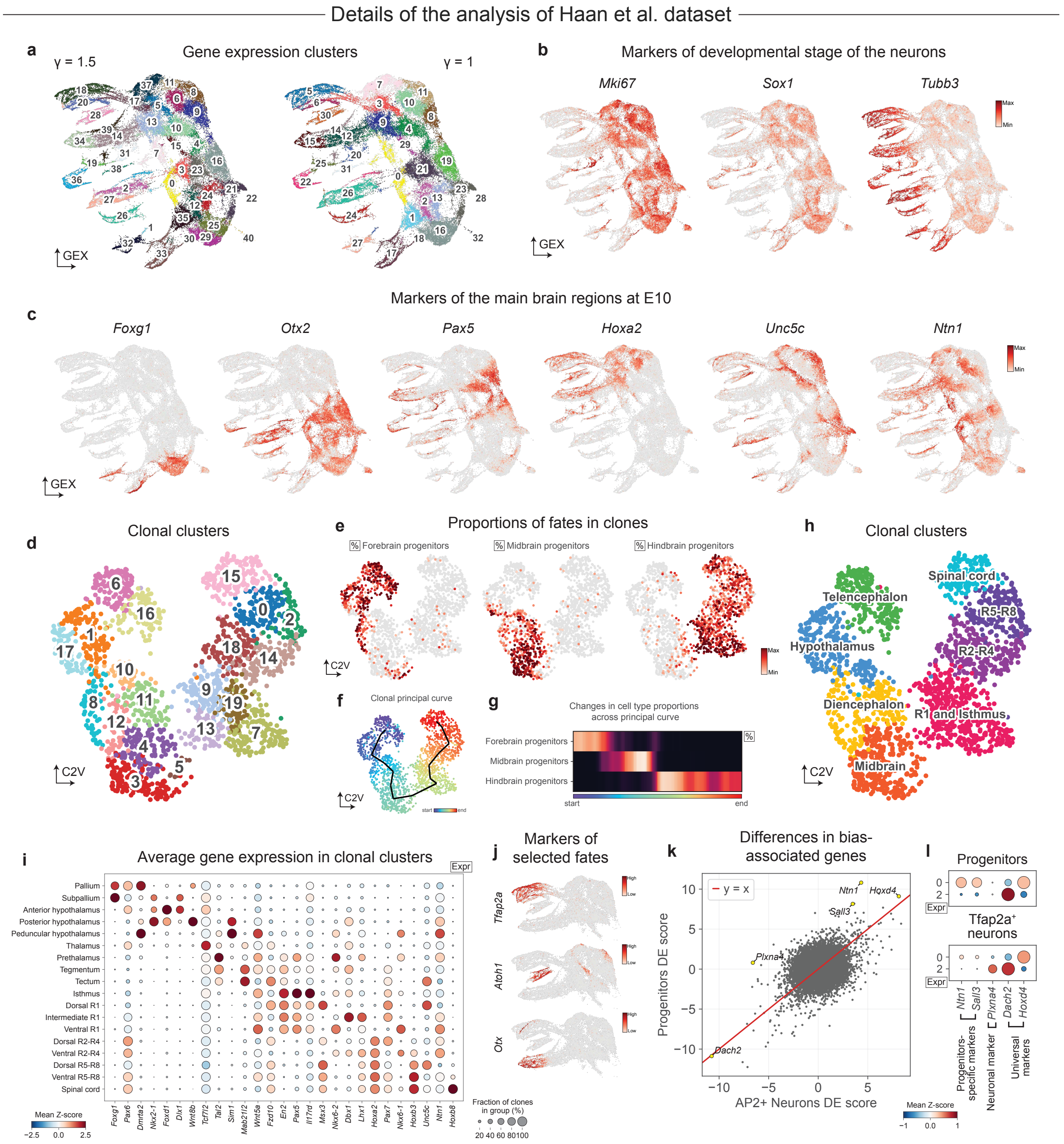


**Supplementary Figure 5**. *Details of the analysis of the Haan et al. dataset*

**a**. Gene expression UMAPs of CNS neurons, colored by Leiden clusters at resolution 1.5 (left) and 1 (right). **b**. Same UMAPs as (a), colored by expression of key markers of neuronal developmental stage. **c**. Same UMAPs as (a), colored by key regional gene expression markers. **d**. clone2vec UMAP, colored by Leiden clusters (resolution = 2). **e**. Same UMAP as (d), colored by the per-clone proportion of different progenitors. **f**. Same UMAP as (d), with the fitted EPG principal curve overlaid. **g**. Heatmap of smoothed per-clone progenitor proportions along the pseudotime ordering derived from the trajectory in (f). **h**. Same UMAP as (d), colored by high-level regional annotation of clones. **i**. Clonal expression dot plot of key regional markers across clonal clusters. **j**. Subset of the gene expression UMAP restricted to hindbrain and spinal cord progenitors and neurons, colored by expression of key markers of neuronal subpopulations. **k**. Scatterplot comparing differential expression scores (Welch's t-statistic for the dorsal vs ventral R5-R8 contrast) between progenitors and *Tfap2a*$^{+}$ neurons. **I**. Clone-level dot plot of selected genes that are specific to *Tfap2a*$^{+}$ neurons, specific to progenitors, or commonly differentially expressed between dorsal and ventral R5-R8.

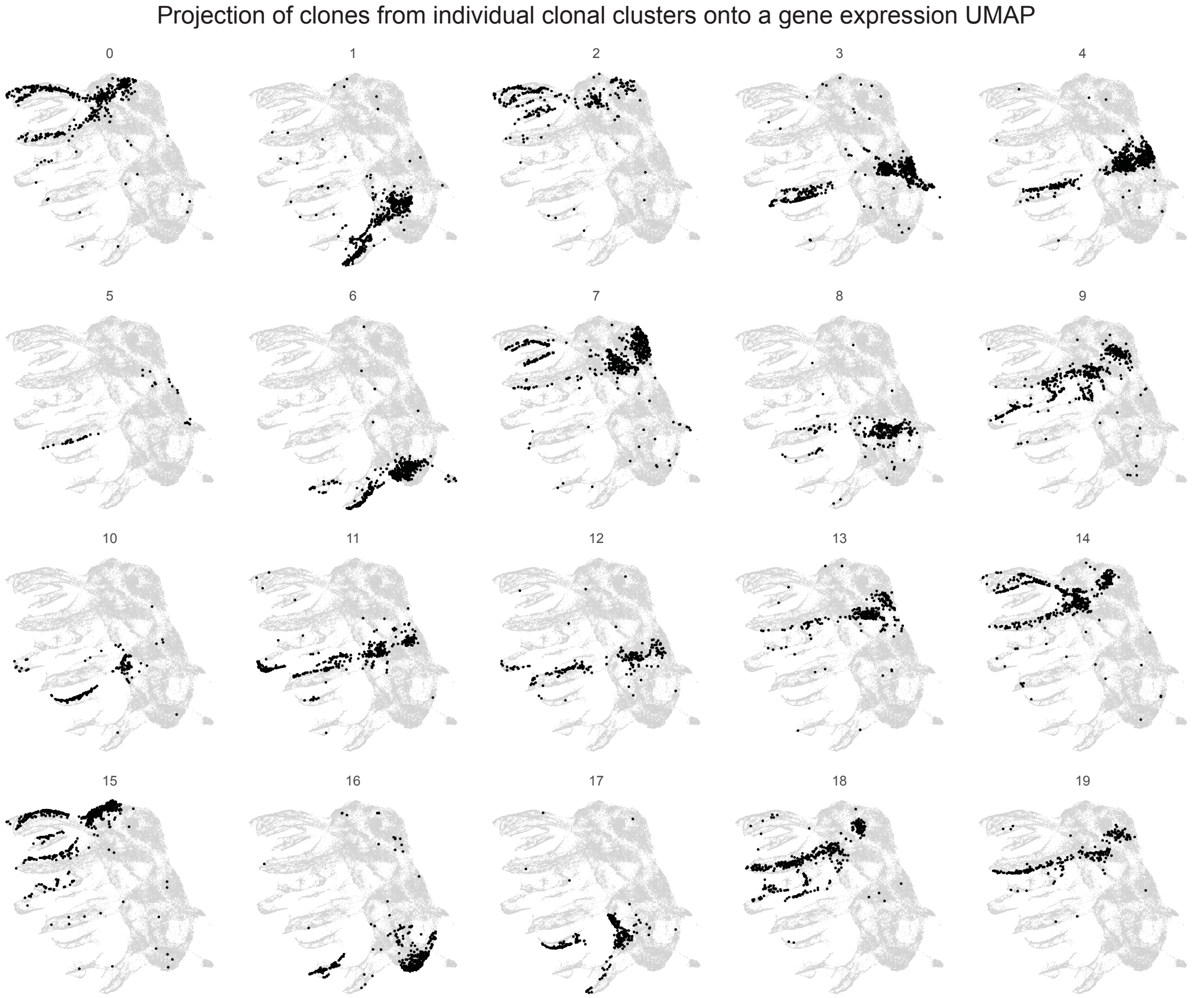


**Supplementary Figure 6**. *Projection of individual clonal clusters (Leiden resolution = 2) onto the gene expression UMAP of CNS neurons from Haan et al.*

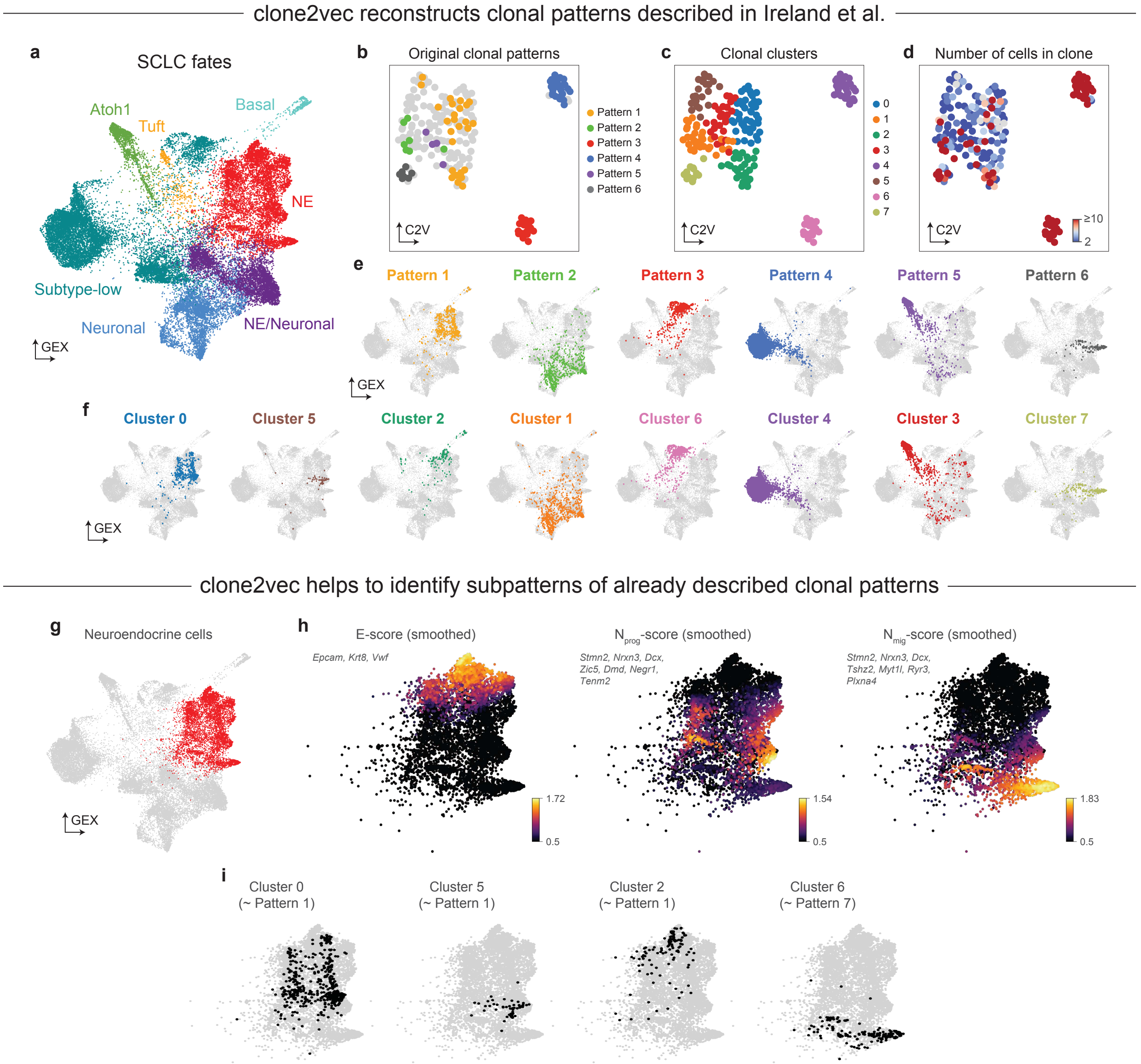


**Supplementary Figure 7**. *Clonal analysis of the SCLC organoid differentiation dataset (Ireland et al.)*

**a**. Gene expression UMAP from the original study, colored by the authors' cell types. **b**. clone2vec UMAP, colored by the authors' clonal patterns; clones not annotated by the authors are shown in grey. **c**. Same UMAP as (b), colored by Leiden clusters. **d**. Same UMAP as (b), colored by clone size. **e**. Projection of clones from each authors' clonal pattern onto the gene expression UMAP. **f**. Projection of clonal clusters from (c) onto the gene expression UMAP. **g**. Same UMAP as (a), with neuroendocrine cells highlighted. **h**. Same UMAP as (a), colored by epithelial (left), neuronal progenitor (center), and migratory neuron (right) gene expression signature scores (signature genes are displayed on the figure). **i**. Projection of selected clonal clusters onto neuroendocrine cells only.

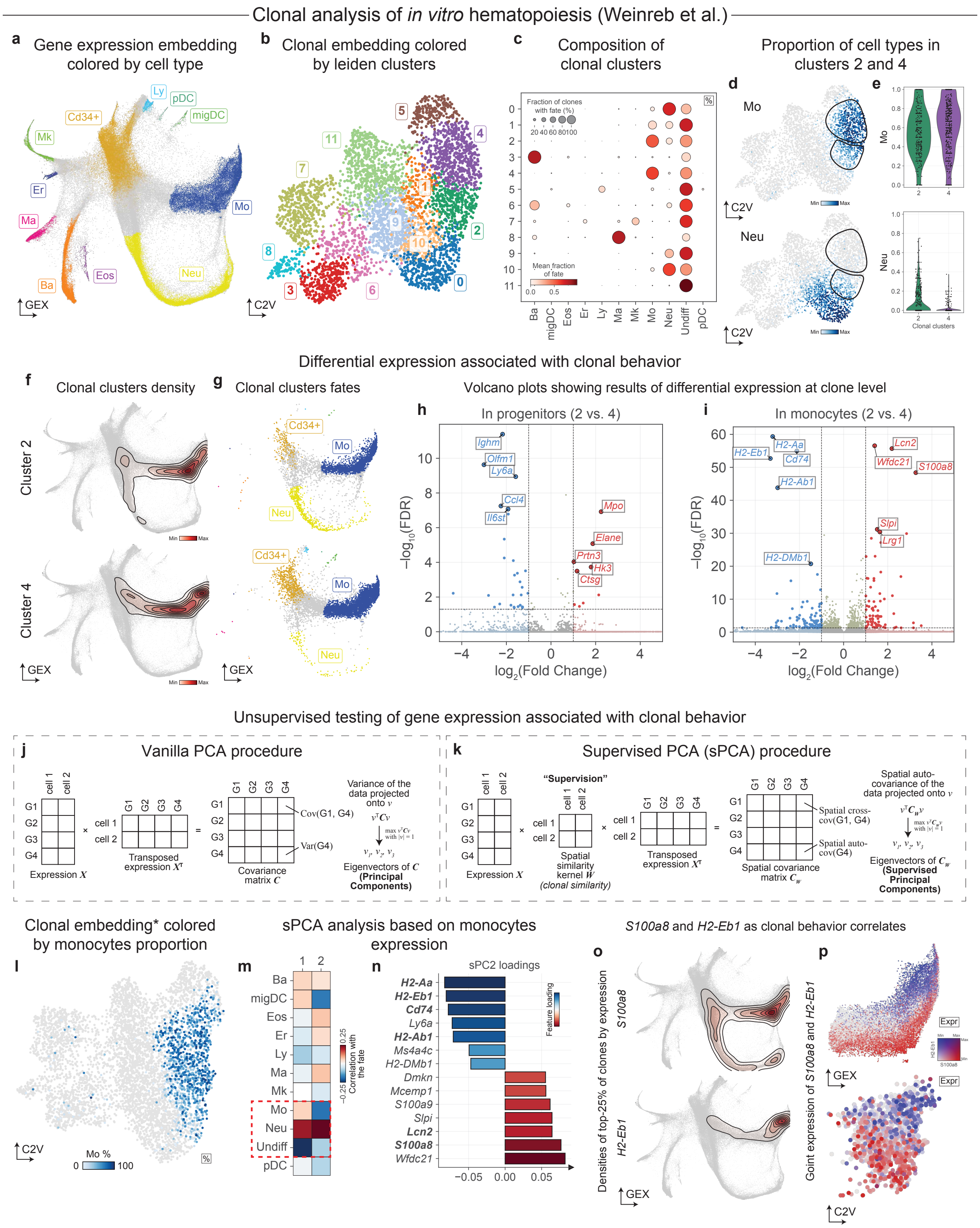


**Supplementary Figure 8**. *Clonal analysis of in vitro hematopoiesis from Weinreb et al.*

**a**. Gene expression UMAP, colored by authors' cell types together with manually identified *Cd34*⁺ progenitors. **b**. clone2vec UMAP, colored by Leiden clusters. **c**. Composition dot plot of clonal clusters (excluding the Cd34+ progenitor population). **d**. Same UMAP as (b), colored by the per-clone proportion of monocytes (top) and neutrophils (bottom); clonal clusters 2 and 4 are outlined. **e**. Violin plots of the per-clone proportions of monocytes (top) and neutrophils (bottom) in clonal clusters 2 and 4. **f**. Gene expression UMAP with KDE of the projections of clonal cluster 2 (top) and 4 (bottom). **g**. Gene expression UMAP showing only cells from clonal clusters 2 (top) and 4 (bottom). **h-i**. Volcano plots of clone-level differential expression (two-sided Welch's t-test with Benjamini-Hochberg correction) between clonal clusters 2 and 4 in progenitors (**h**) and monocytes (**i**). **j**. Schematic of standard PCA. **k**. Schematic of supervised PCA (sPCA). **l**. clone2vec UMAP (built with monocytes masked, as described in Methods), colored by the per-clone proportion of monocytes. **m**. Heatmap of Pearson correlations between sPC coordinates and the proportions of different cell types. **n**. Bar plot of loadings of the top 7 genes with positive and negative loadings on sPC2. **o**. Gene expression UMAP with KDE of the top 25% of clones by mean expression of *S100a8* (top) and *H2-Eb1* (bottom). **p**. Gene expression (top) and clone2vec (bottom) UMAPs bi-colored by expression of *H2-Eb1* (blue) and *S100a8* (red).

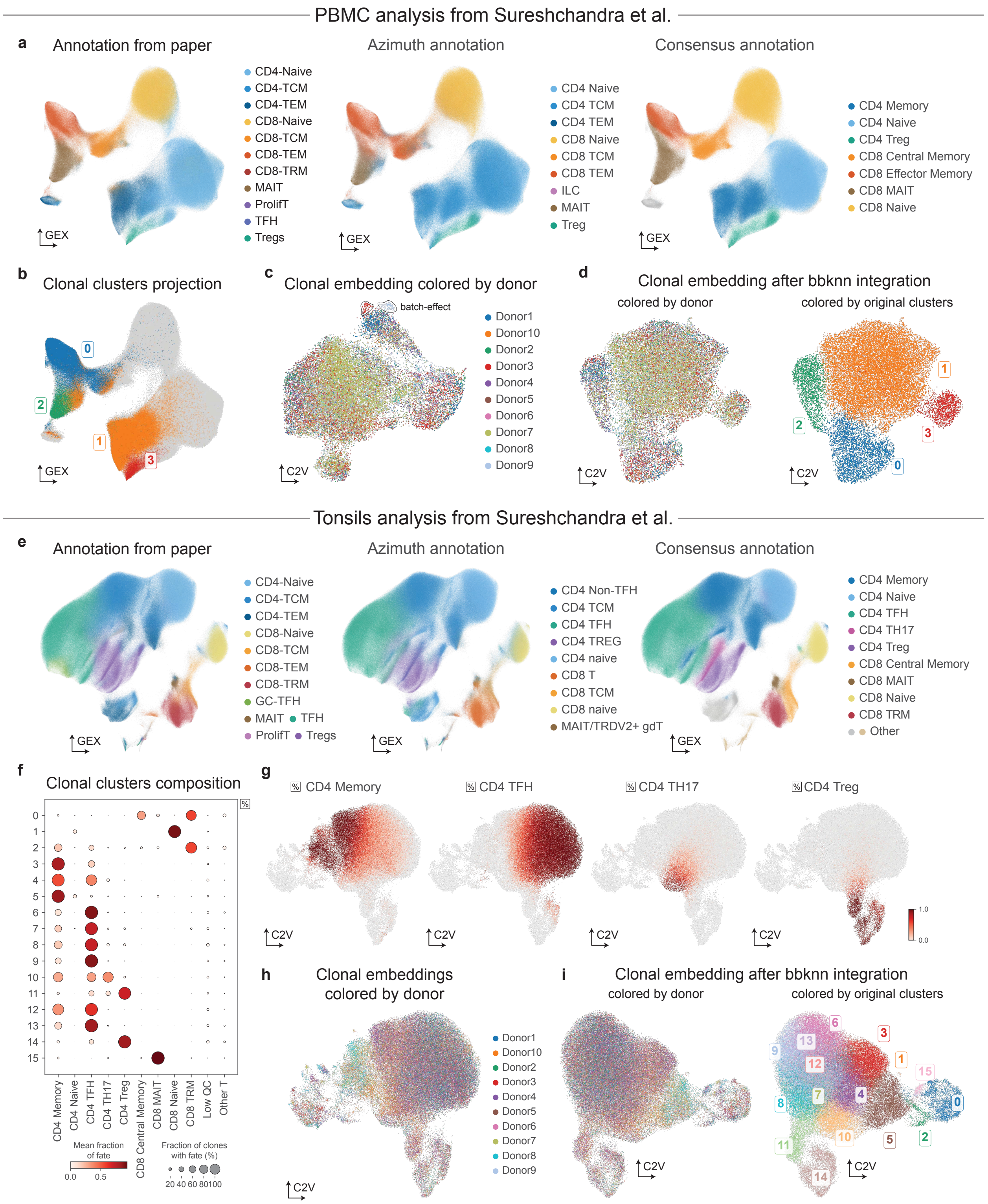


**Supplementary Figure 9**. *Details of the analysis of PBMC and tonsil datasets from Sureshchandra et al.*

**a**. Gene expression UMAPs of PBMC T cells, colored by authors' annotation (left), Azimuth label-transfer annotation (middle), and consensus annotation (right). **b**. Same UMAP as (a), colored by clonal cluster projection. **c**. clone2vec

UMAP, colored by clone donor. **d**. bbknn-corrected clone2vec UMAP, colored by clone donor (left) and uncorrected clusters (right), showing that batch effect does not influence clustering at this resolution. **e**. Gene expression UMAPs of tonsil T cells, colored by authors' annotation (left), Azimuth label-transfer annotation (middle), and consensus annotation (right). **f**. Composition dot plot for each clonal cluster. **g**. clone2vec UMAP, colored by the per-clone proportion of CD4 fates (memory, TFH, Th17, and Treg). **h**. clone2vec UMAP, colored by clone donor. **i**. bbknn-corrected clone2vec UMAP, colored by clone donor (left) and uncorrected clusters (right).

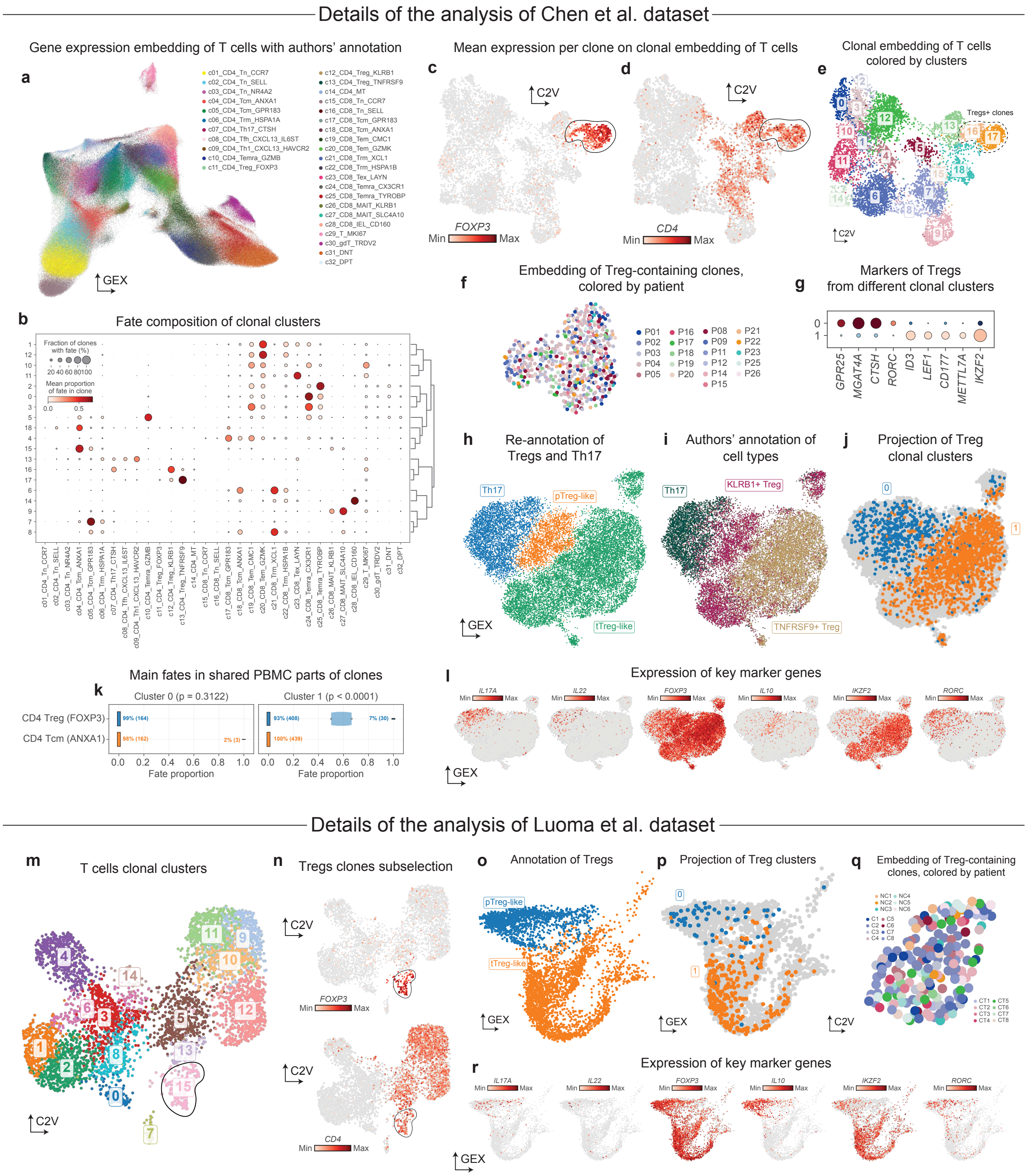


**Supplementary Figure 10**. *Details of the analysis of colon T cell datasets from Chen et al. and Luoma et al.*

(**a-l**) Chen et al. dataset. **a**. Gene expression UMAP of T cells from Chen et al., colored by the authors' cell types. **b**. Composition dot plot for each clonal cluster. **c-d**. clone2vec UMAP of all T cells, colored by per-clone mean expression of *FOXP3* (**c**) and *CD4* (**d**); Treg-containing clones are outlined. **e**. clone2vec UMAP of all T cells, colored by Leiden clusters; Treg-containing clones (clusters 16 and 17) are outlined with a dotted line. **f**. clone2vec UMAP of Treg-containing clones only, colored by clone donor. **g**. Clonal expression dot plot of selected Treg markers across the two

Treg-containing clonal clusters (0 and 1). **h-i**. Gene expression UMAP restricted to Tregs and Th17 cells, colored by manual re-annotation into Th17, pTreg-like and tTreg-like populations (**h**) and by the authors' cell types (**i**). **j**. Projection of Treg-containing clonal clusters 0 and 1 onto the embedding from (h). **k**. Modified violin plots (with bars at 0 and 1 sized according to the fraction of values equal to those numbers) of CD4 Treg (*FOXP3*$^{+}$) and CD4 Tcm (*ANXA1*$^{+}$) fate proportions in matched PBMCs for clones shared with Treg-containing cluster 0 (left, $p = 0.3122$) and cluster 1 (right, $p < 0.0001$). P-values: two-sided Mann-Whitney U-test, without multiple testing correction; the trend is consistent with Fig. 3h, although the cluster 0 comparison does not reach significance. **l**. Gene expression UMAP from (h), colored by expression of key Treg markers.

(**m-r**) Luoma et al. dataset. **m**. clone2vec UMAP of all T cell clones from Luoma et al., colored by Leiden clusters; the Treg-containing cluster (15) is outlined. **n**. Same UMAP as (m), colored by per-clone mean expression of *FOXP3* (top) and *CD4* (bottom); Treg-containing clones are outlined. **o**. Gene expression UMAP restricted to Tregs, colored by manually identified pTreg-like and tTreg-like subpopulations. **p**. Projection of Treg-containing clonal clusters 0 and 1 onto the UMAP from (o); this projection guided the cluster annotation in (o). **q**. clone2vec UMAP of Treg-containing clones, colored by clone donor. **r**. Same gene expression UMAP as (o), colored by expression of key Treg markers.

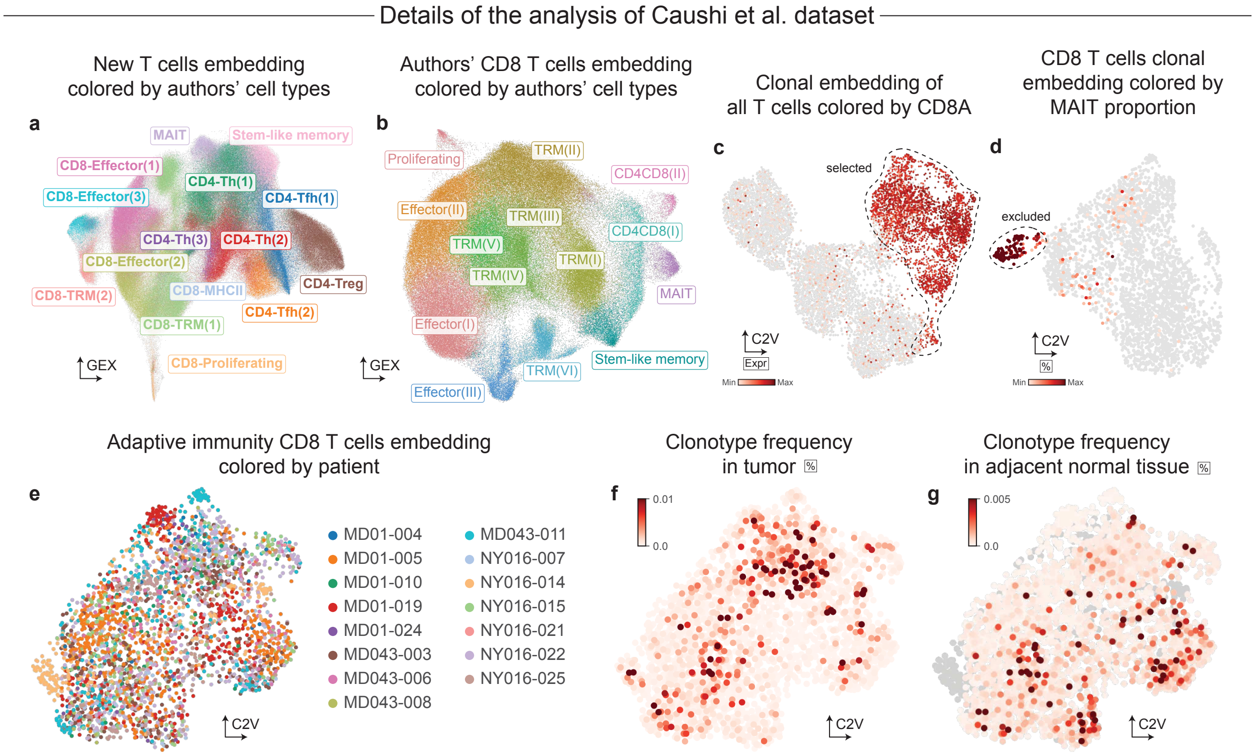


**Supplementary Figure 11**. *Details of the analysis of T cells from NSCLC patients (Caushi et al.)* **a**. De novo gene expression UMAP of all T cells, colored by authors' cell types. **b**. Authors' gene expression UMAP of CD8 T cells, colored by authors' annotation. **c**. clone2vec UMAP of all T cells, colored by per-clone mean expression of CD8A; clones included in downstream analysis are outlined. **d**. clone2vec UMAP of CD8 T cells, colored by the per-clone proportion of MAIT cells; MAIT-high clonal clusters are outlined. **e**. clone2vec UMAP of adaptive CD8 T cells, colored by clone donor. **f-g**. Same UMAP as (e), colored by clonotype frequency in tumor (**f**) and in adjacent normal tissue (**g**).

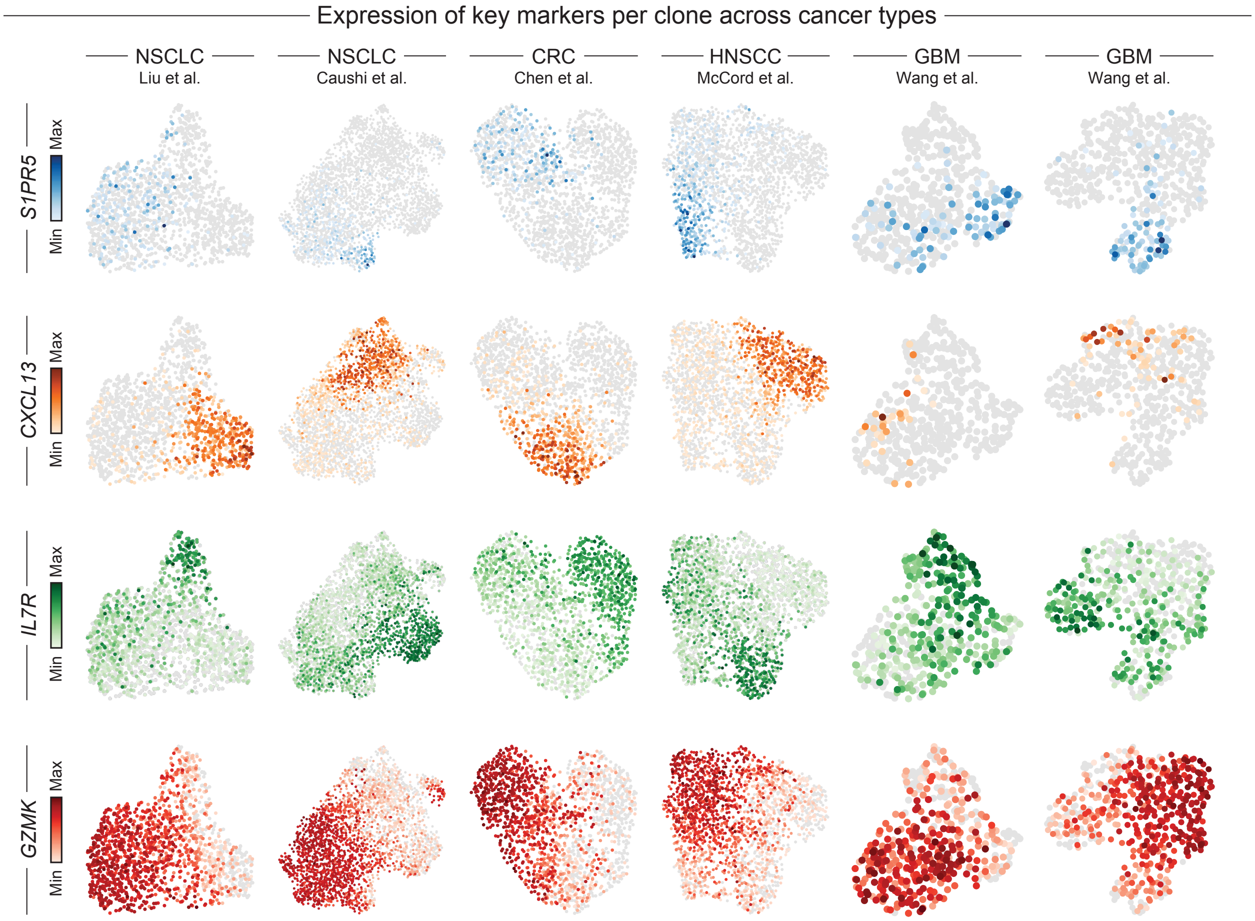


**Supplementary Figure 12**. *Per-archetype marker expression across cancer types*
clone2vec UMAPs of adaptive CD8 T cell lineages across lung cancer (Liu et al., column 1; Caushi et al., column 2), colorectal cancer (Chen et al., column 3), head and neck cancer (McCord et al., column 4), and glioblastoma (Wang et al., multimodal cohort, column 5; Wang et al., scRNA-seq cohort, column 6), colored by expression of *S1PR5* (archetype 0 marker, row 1), *CXCL13* (archetype 1 marker, row 2), *IL7R* (archetype 2 marker, row 3), and *GZMK* (archetype 3 marker, row 4).

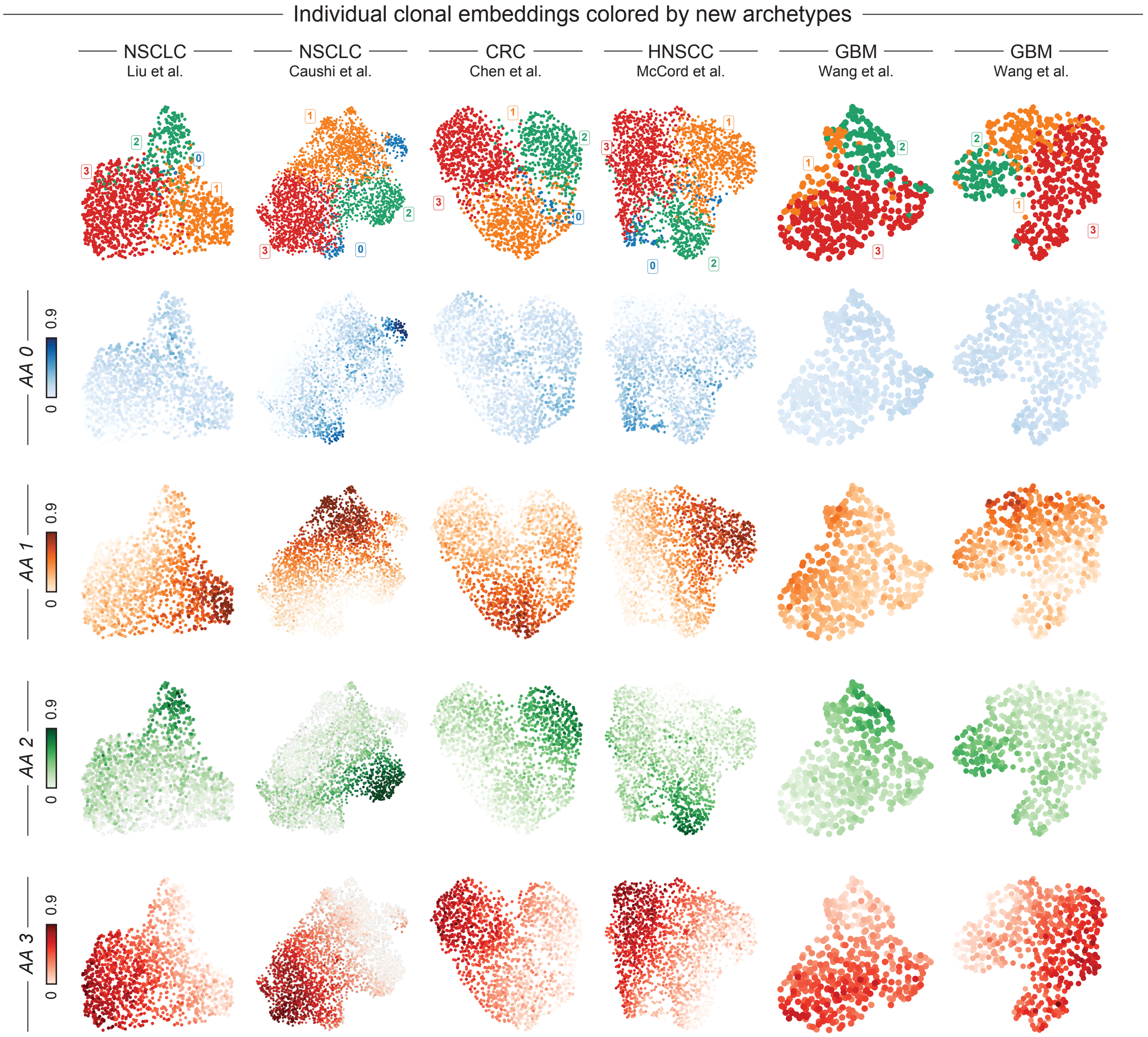


**Supplementary Figure 13**. *Integrated archetype weights across cancer types*
clone2vec UMAPs of adaptive CD8 T cell lineages across lung cancer (Liu et al., column 1; Caushi et al., column 2), colorectal cancer (Chen et al., column 3), head and neck cancer (McCord et al., column 4), and glioblastoma (Wang et al., multimodal cohort, column 5; Wang et al., scRNA-seq cohort, column 6), colored by the weight of integrated archetype 0 (row 1), 1 (row 2), 2 (row 3), and 3 (row 4).

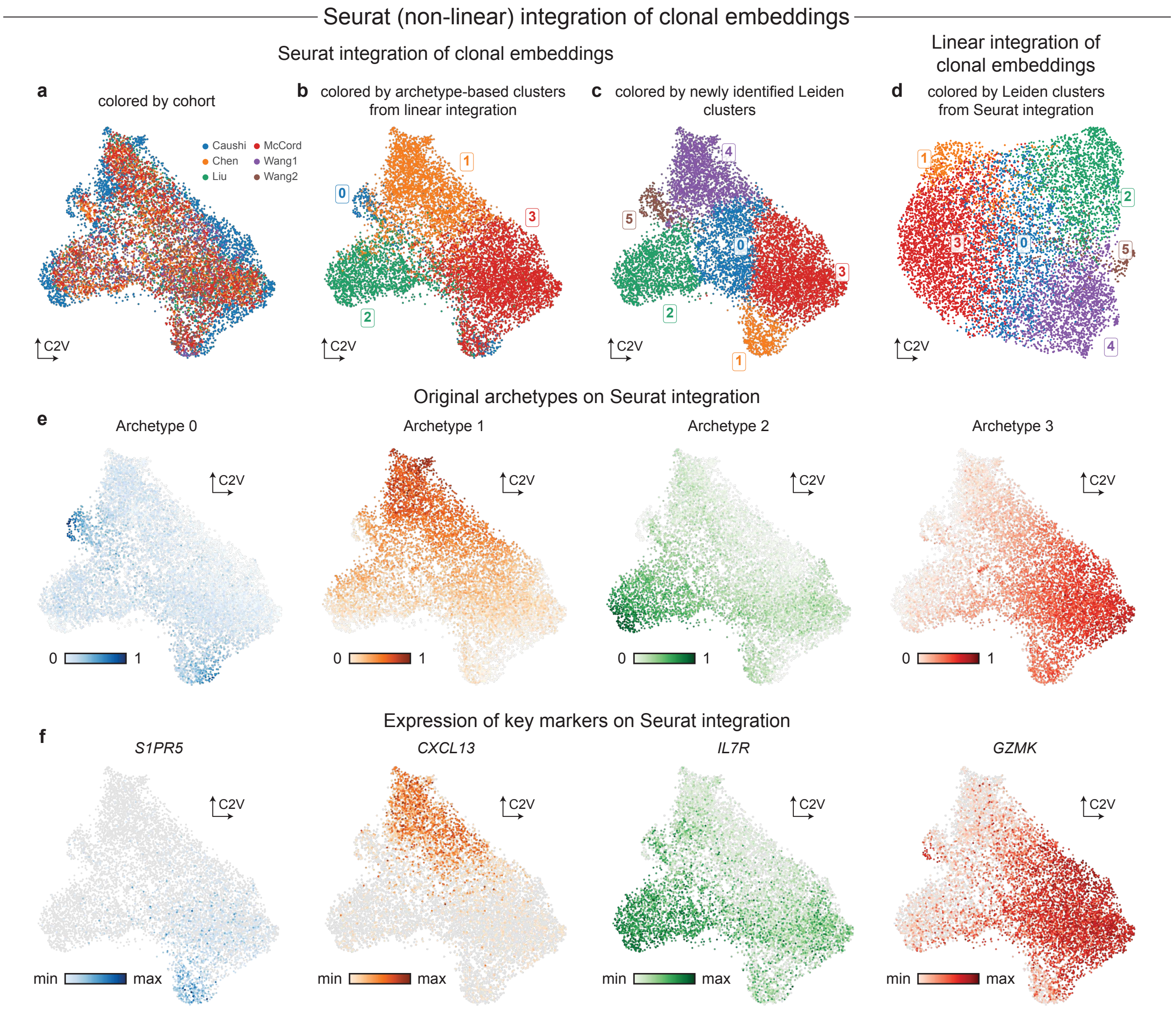


**Supplementary Figure 14**. *Comparison of the proposed integration of clonal embeddings to a Seurat-based approach*

**a**. UMAP of Seurat-integrated clone2vec embeddings, colored by cohort of origin. **b**. Same UMAP as (a), colored by archetype-based clusters from our integration. **c**. Same UMAP as (a), colored by Leiden clusters from the Seurat integration. **d**. UMAP of our clone2vec integration, colored by Seurat clusters from (c). **e**. Same UMAP as (a), colored by the weights of integrated archetypes 0-3 (same order as in Fig. 5h). **f**. Same UMAP as (a), colored by the key archetype markers from Fig. 5c (*S1PR5*, *CXCL13*, *IL7R*, *GZMK*; archetypes 0-3 respectively).

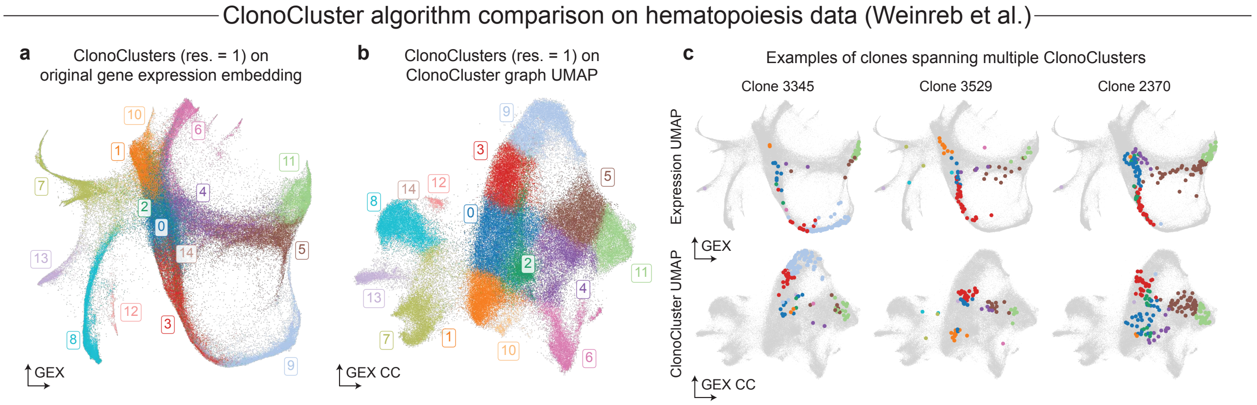


**Supplementary Figure 15**. *Differences in purpose between clone2vec and ClonoCluster*

**a**. Gene expression UMAP of cells from Weinreb et al., colored by ClonoClusters (resolution = 1). **b**. Cell-level UMAP constructed from the ClonoCluster graph, colored by clusters from (a). **c**. Three example clones whose cells are split across multiple ClonoClusters, shown on the gene expression UMAP (top row) and on the ClonoCluster UMAP (bottom row), illustrating that ClonoCluster operates at the cell level – cells from a single clone can be assigned to different clusters – whereas clone2vec assigns a single representation to each clone.
