## Supplementary Note for "Clonal embeddings allow exploratory analysis of lineage-resolved single-cell data"

### Supplementary Note: Formal Description of `clone2vec`

#### 1 Motivation

Quantifying variation in the distribution of clonal fates is useful for identifying stereotypical fate biases and enables downstream analyses such as the detection of developmental regulators of fate. High-throughput lineage tracing makes it possible to recover the progeny distributions of thousands of progenitor cells across cell types, but suffers from both false negatives (*dropouts*) and false positives, so a robust metric on the space of clones is needed.

Clones are traditionally represented as vectors of compositions across gene-expression clusters (cell types). However, within the same cell type, clones with distinct behaviors often carry characteristic expression signatures of those behaviors, making them more similar to each other than to clones with different behaviors in the same cell type [1]. Together with the uncertainty inherent in cell-type identification [2], this motivates a nearest-neighbor graph representation of cells instead. Here we propose `clone2vec`, a method for computing distances between clones based on the similarity of their co-occurrences in such a graph.

#### 2 Formal description

Throughout this note we use $i, j$ to index individual cells and $u, v$ to index individual clones.

Let $\mathcal{X} = \{x_1, \ldots, x_N\}$ be the set of cells with clonal barcodes, where $x_i \in \mathbb{R}^d$ are the coordinates of the $i$-th cell in the expression space (in practice, a PCA or PCA-like embedding such as Harmony or scVI latent space). Let $C = \{C_1, \ldots, C_M\}$ be a partition of $\mathcal{X}$ into $M$ disjoint sets (*clones*), and let $\mathbf{I} : \mathcal{X} \to \{1, \ldots, M\}$ be the clonal identity function, so that $\mathbf{I}(x_i) = c_i$ is the index of the clone containing the $i$-th cell ($x_i \in C_{c_i}$).

For each cell $x_i$, the expression neighborhood (or expression context) $\mathcal{N}(x_i)$ is the set of its $k$ nearest neighbors under the Euclidean (or any other useful) distance in the expression space. Our goal is to learn a low-dimensional embedding $\mathbf{E} \in \mathbb{R}^{M \times h}$ with $h \ll M$, in which each clone $C_u$ is represented by a vector $\mathbf{e}_u \in \mathbb{R}^h$ and clones with similar neighborhood compositions lie close together.

We adopt a Skip-Gram model [3] in which the target is a cell $x_i$ with clonal identity $c_i$, and the context is the multiset of clonal identities of its expression neighbors, $\{c_j : x_j \in \mathcal{N}(x_i)\}$ (multiplicity matters because several neighbors may belong to the same clone). Let $\mathbf{W} \in \mathbb{R}^{M \times h}$ and $\mathbf{W}' \in \mathbb{R}^{h \times M}$ be the input-to-hidden and hidden-to-output weight matrices. We write $\mathbf{w}_u \in \mathbb{R}^h$ for the $u$-th row of $\mathbf{W}$ (the *embedding vector* of clone $C_u$) and $\mathbf{w}'_v \in \mathbb{R}^h$ for the $v$-th column of $\mathbf{W}'$

(the *projection vector* of clone $C_v$). Every clone plays two roles in the model: sometimes it is the *target* clone whose neighborhood is being predicted, and sometimes it is a *neighbor* clone being predicted in someone else's neighborhood. The model assigns each clone one vector for each role: $\mathbf{w}_u$ is used when clone $u$ is the target, and $\mathbf{w}'_v$ is used when clone $v$ is being predicted as a neighbor. Only the target-role vectors $\mathbf{w}_u$ are used downstream as the final clonal embedding.

Given the clonal identity $c$ of a target cell, the probability that a neighbor has clonal identity $c'$ is modeled as

$$P(c' \mid c) = \frac{\exp(\mathbf{w}_c \cdot \mathbf{w}'_{c'})}{\sum_{v=1}^{M} \exp(\mathbf{w}_c \cdot \mathbf{w}'_v)}. \tag{1}$$

The training log-likelihood aggregates this over all observed target–neighbor pairs:

$$\ell = \sum_{i=1}^{N} \sum_{x_j \in \mathcal{N}(x_i)} \log P(c_j \mid c_i). \tag{2}$$

The model is trained by minimizing $-\ell$, and after convergence the rows of $\mathbf{W}$ form the latent clonal embedding $\mathbf{E}$.

### 3 Properties of the clonal embedding

Consider the distribution over directed (target, neighbor) pairs induced by sampling a uniformly random edge of the $k$-nearest-neighbor graph. Let $p(c, c')$ denote the probability that such a random pair has target in clone $c$ and neighbor in clone $c'$. The corresponding target and neighbor marginals are

$$p_{\mathrm{t}}(c) = \sum_{c'} p(c, c'), \qquad p_{\mathrm{n}}(c) = \sum_{c'} p(c', c),$$

that is, $p_{\mathrm{t}}(c)$ is obtained by summing out the neighbor slot and $p_{\mathrm{n}}(c)$ by summing out the target slot. In general $p_{\mathrm{t}} \neq p_{\mathrm{n}}$, since the $k$-NN graph is directed and its in-degree distribution is not uniform. Let $p(c' \mid c) = p(c, c')/p_{\mathrm{t}}(c)$ denote the conditional probability that a neighbor belongs to clone $c'$ given that the target is in clone $c$.

A Skip-Gram model with sufficient capacity approximates these empirical conditionals at convergence [4]. The statements in this section are formulated in the population limit where $P(c' \mid c) = p(c' \mid c)$ exactly; in practice — with finite data and a low-rank ($h \ll M$) embedding — they hold approximately. The identities below take logarithms of $p(c' \mid c)$ and therefore assume $p(c' \mid c) > 0$ for all $c, c'$ of interest. We also write

$$Z_c = \sum_{v=1}^{M} \exp(\mathbf{w}_c \cdot \mathbf{w}'_v) \tag{3}$$

for the softmax normalization constant of (1).

##### 3.1 The PMI factorization

**What is PMI?** The *pointwise mutual information* (PMI) of two events $A$ and $B$ measures how much more or less often they co-occur than independence would predict:

$$\mathrm{PMI}(A, B) = \log \frac{p(A, B)}{p(A)\, p(B)}. \tag{4}$$

A value of zero means $A$ and $B$ co-occur exactly at the rate expected under independence; positive values indicate enrichment, negative values indicate depletion. In our setting, the two events are the identities of a target cell and one of its expression neighbors. Thus $\mathrm{PMI}(u, v)$ measures how much more often clone $v$ appears in the expression neighborhoods of cells from clone $u$ than its marginal frequency would predict, i.e.

$$\mathrm{PMI}(u, v) = \log \frac{p(u, v)}{p_{\mathrm{t}}(u)\, p_{\mathrm{n}}(v)}.$$

Because the $k$-nearest-neighbor graph is directed, $\mathrm{PMI}(u, v) \neq \mathrm{PMI}(v, u)$ in general.

**Theorem 1** (PMI factorization)**.** *At convergence of the Skip-Gram objective,*

$$\mathbf{w}_u \cdot \mathbf{w}'_v = \mathrm{PMI}(u, v) + \log p_{\mathrm{n}}(v) + \log Z_u. \tag{5}$$

*Proof.* From $P(v \mid u) = p(v \mid u) = p(u, v)/p_{\mathrm{t}}(u)$ and the softmax (1),

$$\exp(\mathbf{w}_u \cdot \mathbf{w}'_v) = Z_u \cdot \frac{p(u, v)}{p_{\mathrm{t}}(u)}.$$

Taking logarithms and adding and subtracting $\log p_{\mathrm{n}}(v)$,

$$\mathbf{w}_u \cdot \mathbf{w}'_v = \log \frac{p(u, v)}{p_{\mathrm{t}}(u)\, p_{\mathrm{n}}(v)} + \log p_{\mathrm{n}}(v) + \log Z_u,$$

which is (5). □

Theorem 1 is the central structural result for `clone2vec`: the matrix of inner products $[\mathbf{w}_u \cdot \mathbf{w}'_v]$ equals the PMI matrix plus additive row terms $\log Z_u$ and column terms $\log p_{\mathrm{n}}(v)$. For later use, we record the equivalent form obtained by expanding $\mathrm{PMI}(u, v)$ in (5):

$$\mathbf{w}_u \cdot \mathbf{w}'_v = \log p(v \mid u) + \log Z_u. \tag{6}$$

##### 3.2 Consequences of PMI: robustness to sparse sampling

The PMI factorization has two immediate implications. First, the embedding geometry encodes conditional co-occurrence probabilities, not the positions or counts of individual cells. Second, each embedding vector is a parameter in a low-rank joint fit over the entire co-occurrence matrix, rather than a summary of one clone's cells in isolation. Together, these properties make `clone2vec` well-suited to the sparse, uneven sampling typical of lineage tracing data, as we develop in the three corollaries and the remark below.

**Corollary 2** (Local structure preservation)**.** *If clones $C_u$ and $C_v$ induce similar conditional distributions, $p(c' \mid u) \approx p(c' \mid v)$ for all $c'$, and the projection vectors $\{\mathbf{w}'_{c'}\}_{c'=1}^{M}$ are affinely full-rank (that is, the differences $\mathbf{w}'_{c'} - \mathbf{w}'_{c''}$ span $\mathbb{R}^h$), then $\mathbf{w}_u \approx \mathbf{w}_v$.*

*Proof.* By (6), applied to $u$ and $v$ and then subtracted,

$$(\mathbf{w}_u - \mathbf{w}_v) \cdot \mathbf{w}'_{c'} = \log p(c' \mid u) - \log p(c' \mid v) + \log Z_u - \log Z_v \approx \log Z_u - \log Z_v,$$

which is constant in $c'$. Subtracting the same identity for a second label $c''$ eliminates the constant:

$$(\mathbf{w}_u - \mathbf{w}_v) \cdot (\mathbf{w}'_{c'} - \mathbf{w}'_{c''}) \approx 0 \qquad \text{for all } c', c''.$$

Under the affine-full-rank assumption, $\{\mathbf{w}'_{c'} - \mathbf{w}'_{c''}\}$ spans $\mathbb{R}^h$, so the only vector orthogonal to all of them is zero; hence $\mathbf{w}_u \approx \mathbf{w}_v$. Substituting back into the first identity gives $Z_u \approx Z_v$. □

**Corollary 3** (Clone-size stability)**.** *In the population limit, two clones with similar conditional distributions receive similar embeddings regardless of their sizes.*

*Proof.* The conditional $p(c' \mid c)$ is a frequency ratio, not a count: it depends on the shape of a clone's neighborhood distribution but not on the absolute number of cells. The conclusion follows from Corollary 2. □

**Corollary 4** (Dropout stability)**.** *Suppose the cells of $C_u$ share similar expression neighborhoods, $\mathcal{N}(x) \approx \mathcal{N}(y)$ for all $x, y \in C_u$ (**neighborhood consistency**). Then a random subsample $C'_u \subset C_u$ of sufficient size induces approximately the same empirical conditional distribution (with high probability) and, by Corollary 2, approximately the same embedding.*

The three corollaries above hold at the Skip-Gram MLE. A related but distinct question is whether the MLE itself is well-estimated when clones are small, and how it compares to sample-based alternatives in that regime. The following remark is informal.

*Remark* (Robustness of the estimator). At the sparse clone sizes typical of lineage tracing, we expect the embedding distance $\|\mathbf{w}_u - \mathbf{w}_v\|$ to be a more stable measure of clonal dissimilarity than direct sample statistics such as MMD or optimal transport. Two design choices contribute.

MMD and optimal transport are nonparametric distances between empirical distributions on the continuous expression manifold $\mathbb{R}^d$. They are designed for a regime in which each sample is represented by enough points to approximate its underlying density, and become unreliable in the sparse regime where clones are represented by two to five cells: the empirical distribution is then too coarse to stand in for the true one, and the computed distance is dominated by sampling noise.

`clone2vec` is built around a structurally different statistical object. Every neighborhood observation is a nonnegative integer count in the $M \times M$ co-occurrence matrix $\mathbf{N}$, and the embedding is a multinomial factorization of $\mathbf{N}$ (see §5). This brings two advantages in the sparse regime:

(i) *Discrete coarse-graining.* The model asks only "how often did a cell of clone $v$ appear next to a cell of clone $u$?". This reduces the estimation problem from a density on $\mathbb{R}^d$ to a categorical co-occurrence over the $M$ clones, which is well-defined even for a handful of cells.

(ii) *Global low-rank pooling*. Every entry of $\mathbf{N}$ contributes to every embedding vector through the shared projection vectors $\mathbf{w}'_v$, and the rank-$h$ constraint forces all embeddings into a common low-dimensional subspace. A small clone whose neighborhoods overlap with better-sampled clones therefore inherits its position from them, rather than relying on its own handful of cells.

Direct sample statistics have neither property: they operate on continuous empirical distributions and pool no information across clones.

##### 3.3 Global structure preservation

**Theorem 5** (Global structure preservation, first-order)**.** *Suppose clone $C_w$ is a convex mixture of clones $C_u$ and $C_v$, i.e. $p(c' \mid w) \approx \alpha\, p(c' \mid u) + (1-\alpha)\, p(c' \mid v)$ for some $\alpha \in [0,1]$ and all $c'$. Then to first order in a linearization of the softmax around the convex combination,*

$$\mathbf{w}_w \approx \alpha\, \mathbf{w}_u + (1-\alpha)\, \mathbf{w}_v.$$

This does not follow directly from Theorem 1, because the PMI factorization involves logarithms of the conditional distributions and log does not commute with convex combinations. For the full proof, see Theorem 2 of [5].

Together, Theorems 1–5 and Corollaries 2–4 show, under the assumptions above, that clonal embeddings preserve both local and global structure, tolerate variation in clone size and sampling depth, and admit a principled information-theoretic interpretation. These properties support downstream analyses ranging from discrete cluster-based summaries to trajectory inference in the continuous clonal space.

#### 4 Model parameters

`clone2vec` has two main hyperparameters: the neighborhood size $k$ and the embedding dimension $h$.

**Neighborhood size $k$.** This parameter sets the effective radius in expression space within which cells are treated as sharing a similar state. Larger datasets with denser sampling of the expression manifold can accommodate larger $k$, which both enlarges the effective training set and improves embedding quality. However, very large $k$ blurs distinct cell states and increases the training cost, which scales as $O(k \cdot h \cdot M \cdot N)$ under the full-softmax objective. Values of $k = 10$–$15$ work well in most datasets.

**Embedding dimension $h$.** This parameter controls the intrinsic dimensionality of the clonal space. It should be large enough to capture the relevant axes of clonal behavior but not so large that the clonal manifold loses smoothness or overfits. Values of $h = 5$ (for simple manifolds with a few clonal behaviors) to $h = 10$ (for more complex ones) work well in practice.

### 5 GLM-PCA formulation and Poisson approximation

#### 5.1 Skip-Gram as multinomial GLM-PCA

The Skip-Gram objective admits an equivalent formulation as generalized linear model PCA (GLM-PCA), also called exponential-family PCA in the earlier literature [6]. Define the clone-by-clone co-occurrence count matrix $\mathbf{N} \in \mathbb{N}^{M\times M}$ by

$$N_{uv} = \sum_{x\in C_u} \left|\{y \in \mathcal{N}(x) : \mathbf{I}(y) = v\}\right|, \tag{7}$$

so that $N_{uv}$ counts how often cells of clone $v$ appear in the expression neighborhoods of cells of clone $u$. The row and column sums recover the target and neighbor marginals up to the total edge count: $\sum_v N_{uv} \propto p_\text{t}(u)$ and $\sum_u N_{uv} \propto p_\text{n}(v)$. Under the Skip-Gram likelihood, each row $\mathbf{N}_{u,\cdot}$ is modeled as a multinomial draw with probabilities

$$\pi_{uv} = \frac{\exp(\mathbf{w}_u \cdot \mathbf{w}'_v)}{\sum_{v'=1}^{M} \exp(\mathbf{w}_u \cdot \mathbf{w}'_{v'})},$$

so maximizing (2) is equivalent to fitting a rank-$h$ factorization of the row-wise log-probabilities of $\mathbf{N}$ under a multinomial observation model — that is, multinomial GLM-PCA applied to $\mathbf{N}$ [6, 7].

#### 5.2 Poisson approximation and `fastglmpca`

The multinomial formulation has a practical drawback: the softmax couples all $M$ output units through the partition constant $Z_u$, so the gradient with respect to $\mathbf{w}_u$ depends on every projection vector. This prevents the optimization from decomposing into independent sub-problems and becomes limiting when $M$ reaches the order of $10^5$, as in large immune-receptor datasets.

A standard way to decouple the factors is the *Poisson trick* [8]: conditional on the row sums, the multinomial likelihood on each row of $\mathbf{N}$ is equivalent (up to a factor depending only on the row sum) to a product of independent Poisson likelihoods on its entries. This motivates the *offset-augmented log-bilinear model*

$$N_{uv} \sim \text{Poisson}(\lambda_{uv}), \qquad \log \lambda_{uv} = \mathbf{w}_u \cdot \mathbf{w}'_v + a_u + b_v, \tag{8}$$

where $a_u$ and $b_v$ absorb row (target-marginal) and column (neighbor-marginal) counts, respectively. This is not exactly equivalent to the multinomial Skip-Gram: the column offsets $b_v$ shift the implied softmax probabilities by a factor of $\exp(b_v)$. It does, however, target the same interaction structure up to row- and column-wise shifts, as made precise in §5.3 below. With $\mathbf{W}'$ and the column offsets $\{b_v\}$ held fixed, updating $\mathbf{W}$ and the row offsets $\{a_u\}$ decomposes into $M$ independent Poisson GLMs (one per row); the dual update in $\mathbf{W}'$ and $\{b_v\}$ decomposes symmetrically across columns.

This structure is exploited by the Alternating Poisson Regression (APR) algorithm in `fastglmpca` [9]. APR alternates between the two half-steps above; each is trivially parallel across rows (or columns) and monotonically improves the log-likelihood, yielding a memory-efficient, multi-core-friendly optimizer that scales to datasets where the multinomial fit is infeasible.

##### 5.3 Relationship between Poisson and multinomial solutions

The Poisson and multinomial fits target the same latent geometry up to row- and column-wise shifts. Comparing (8) with the PMI factorization (5),

$$\mathbf{w}_u \cdot \mathbf{w}'_v \;=\; \log \lambda_{uv} - a_u - b_v \;\approx\; \mathrm{PMI}(u, v) + \mathrm{const},$$

once $a_u$ absorbs the row-marginal contribution (including $\log Z_u$) and $b_v$ absorbs the neighbor marginal $\log p_{\mathrm{n}}(v)$. The equivalence of the interaction term under row- and column-wise offsets is a classical result in the log-linear modeling of contingency tables [8], and ensures that the Poisson fit recovers the same low-rank interaction structure, and hence the same PMI-type dot-product geometry, that underlies Theorem 1. Our numerical experiments (main text, Figs. S1–S3) confirm that the Poisson and multinomial embeddings agree qualitatively, while the Poisson variant is substantially faster to fit.

We use multinomial Skip-Gram as the default, since it matches the compositional structure of neighborhood counts exactly. When the number of clones makes the multinomial fit prohibitive, we use Poisson GLM-PCA via `fastglmpca` with offsets as a fast alternative that recovers the same qualitative clonal geometry.

#### 6 Comparison to the CBOW model

Two standard architectures are used for word-embedding models: Skip-Gram and Continuous Bag-of-Words (CBOW) [3]. Skip-Gram predicts each context element independently from the target, so every target–context pair contributes its own gradient update. CBOW instead predicts the target from the average of its context vectors and therefore updates vectors based on a collective, averaged signal. In the clonal setting, CBOW replaces the target clone embedding in (1) with the average of the embeddings of its context cells.

The two architectures induce qualitatively different geometries. Skip-Gram's pairwise updates reinforce strong local associations and tend to produce more pronounced clustering, whereas CBOW's averaging smooths over individual pairs and yields a more homogeneous latent space [10]. Because our goal is to resolve distinct clonal behaviors rather than to assign average-case labels, we adopt the Skip-Gram formulation throughout.

[9] Eric Weine, Peter Carbonetto, and Matthew Stephens. "Accelerated dimensionality reduction of single-cell RNA sequencing data with fastglmpca". en. In: *Bioinformatics* 40.8 (Aug. 2024).

[10] Xin Rong. *word2vec Parameter Learning Explained*. 2016. arXiv: 1411.2738 [cs.CL]. URL: https://arxiv.org/abs/1411.2738.
